# Sst+ GPi output neurons provide direct feedback to key nodes of the basal ganglia and drive behavioral flexibility

**DOI:** 10.1101/2022.03.16.484460

**Authors:** M. Weglage, S. Ährlund-Richter, A. Contestabile, V. Skara, J. Fuzik, I. Santos, I. Lazaridis, K. Meletis

**Affiliations:** Department of Neuroscience, Karolinska Institutet, Stockholm, Sweden; Department of Organismic and Evolutionary Biology and Center for Brain Science, Harvard University, Cambridge, MA, USA; Picower Institute of Learning and Memory, Massachusetts Institute of Technology, Cambridge, MA, USA; McGovern Institute, Massachusetts Institute of Technology, Cambridge, MA, USA

## Abstract

The internal globus pallidus (GPi) is a basal ganglia output nucleus with separate projections to the thalamus and the lateral habenula (LHb). Here, we show a GPi subtype with projections to LHb (GPi-LHb), genetically defined based on glutamate/GABA co-transmission and somatostatin (Sst) expression, also projects back to key nodes in the basal ganglia: the external globus pallidus (GPe), the striatal striosomes, and dopamine neurons in the substantia nigra. We found that the Sst+ GPi population showed strong movement and direction-specific selectivity in a goal-directed choice task, but not during self-paced exploration, and did not signal prediction errors or outcome modulation. During goal-directed behavior, the Sst+ GPi activity slowly evolved with learning of correct choice actions and genetic silencing disrupted the ability to update choice behavior following a task rule reversal. In summary, we have found that Sst+ GPi neurons establish a wide feedback network in the basal ganglia and drive behavioral flexibility.

## Introduction

The basal ganglia are important for the generation of voluntary movements, decision-making and motivated behaviors^1,2^. The selection of specific actions, and the process of updating the value of chosen actions and strategies, ultimately produces motivated behaviors that optimize gain from the environment. Changes in the environmental conditions, such as resource availability (e.g. food), require animals to dynamically update their actions to survive. From this perspective, the choice to continue performing the same rewarded action (exploitation), versus the exploration of alternative options, is a critical computation. The basal ganglia and the habenula form an evolutionary conserved circuit optimized to support dynamic decision-making and choice updating^3^. The inability to adapt and explore (i.e., behavioral inflexibility) is manifested as habitual or compulsive actions^4^, a condition associated with psychiatric disorders in humans and considerable cost in terms of quality of life^5^.

The internal segment of the globus pallidus (GPi) and the substantia nigra pars reticulata (SNr) are the output nuclei of the basal ganglia, which in the canonical basal ganglia model regulate behavior through projections to thalamus and other downstream motor centers^6,7^. The mouse GPi (also called entopeduncular nucleus) can be further subdivided into two main output pathways: i) the parvalbumin-expressing (Pv+) GPi neurons that project to thalamus and brainstem^8^ and ii) the somatostatin-expressing (Sst+) GPi neurons that project to the lateral habenula (LHb)^9–12^. The Sst+ GPi-LHb neurons co-release GABA and glutamate in LHb^13–15^ and have been suggested to carry information related to the outcomes of actions^16^, although optogenetic activation of the Sst+ GPi-LHb neurons does not modulate action value^13^. Thus, the GPi-LHb pathway holds a unique neuroanatomical position to integrate motor and non-motor information from the basal ganglia to guide and update motivated behavior.

Here, we used genetic targeting of the GPi-LHb neurons, based on their unique molecular profile: the co-expression of the transporters Vglut2 and Vgat (i.e., based on their glutamate/GABA co-transmission profile), or expression of the defining cellular marker Sst^13,15^. We found that visualization of the brain-wide projections of GPi-LHb neurons revealed an unexpected organization: in addition to the well-known axonal projection to LHb, GPi neurons also send projections to the external segment of the globus pallidus (GPe), the striatal striosomes, and the substantia nigra pars compacta (SNc) - three key nodes of the basal ganglia. This organization suggested that GPi-LHb neurons could shape motivated behaviors by calibrating the activity of functional nodes in the basal ganglia circuitry. We therefore recorded the activity of Sst+ GPi-LHb neurons in a serial reversal task using fiber photometry to define their role during motivated behaviors. These recordings showed no correlation of GPi-LHb pathway activity and reward prediction errors - instead, the Sst+ GPi-LHb activity was strongly modulated during motor-related events, showing activation specifically during vigorous movements and ipsiversive turning. Functional perturbation experiments, using cell-type specific tetanus toxin-mediated silencing demonstrated that Sst+ GPi-LHb neurons are critical for behavioral adaptation following reversal after overtraining in a single reversal maze-based choice task. In summary, we propose that a neuromodulatory feedback signal from Sst+ GPi neurons onto key nodes of the basal ganglia is central to flexible behavior by shifting the network away from well-established and reinforced choices to drive exploration of alternative choices.

## Results

### GPi-LHb neurons have feedback connections to key nodes of the basal ganglia

GPi output neurons that project to the LHb (GPi-LHb neurons) can be defined by their GABA and glutamate co-transmission profile and their specific expression of Sst^13,15^. We mapped the brain-wide projection patterns of GPi-LHb neurons, employing three complementary cell-type specific adeno-associated virus (AAV) genetic labeling strategies. First, we used Vglut2-Cre/Vgat-Flp transgenic mice to selectively target GPi-LHb neurons that co-transmit GABA and glutamate with injection into the LHb of a retrogradely transported AAV (AAVrg) that induced Cre- and Flp-dependent expression of ChR2-EYFP (AAVrg-Con/Fon-ChR2-EYFP; experimental strategy referred to as Vglut2/Vgat^GPi-LHb^, N = 7 mice, **Figure 1A**). As expected, this strategy resulted in genetic labeling of GPi neurons with a dense projection to the LHb (**Figure 1B**). In addition to labeling the known GPi-LHb pathway, we found extensive and dense collateral projections from the labeled Vglut2/Vgat^GPi-LHb^ neurons to other subcortical and cortical brain regions, most prominently targeting the GPe, the caudate putamen (CPu), the SNc, and the prefrontal cortex (PFC; **Figure 1C**, **Supplementary Figure 1A**, **Supplementary Figure 2A-J**). We confirmed that the Vglut2/Vgat^GPi-LHb^ axon terminals targeted the GPe using the parvalbumin (Pv) marker expression (**Figure 1D**). The Vglut2/Vgat^GPi-LHb^ collaterals extended further into the CPu, displaying distinct patterns of concentrated axonal arborizations primarily in the dorsal CPu. Due to the discretely organized innervation pattern, we investigated whether the labeled axons preferentially targeted the mu-opioid receptor-expressing (i.e., Oprm1, MOR) striosomes in CPu^17^. Indeed, we found that the axonal arborization of GPi-LHb neurons was limited to MOR+ dense striosomes (**Figure 1E**, **Supplementary Figure 2I-J**). The Vglut2/Vgat^GPi-LHb^ projections were also found in posterior subcortical regions, with the densest projections in the SNc, confirmed by tyrosine hydroxylase (TH) immunolabelling (**Figure 1F**). In addition, we observed axonal labeling in the substantia innominata (SI), but further analysis suggested that this axonal labeling could result from a different population of Vglut2/Vgat-expressing SI neurons (**Supplementary Figure 1B**). We found that the Vglut2/Vgat^GPi-LHb^ strategy also labeled some cell bodies in the ventral tegmental area (VTA) and dorsal raphe nucleus (DRN, **Supplementary Figure 1C**) consistent with prior findings^14,18,19^.

**Figure 1.**
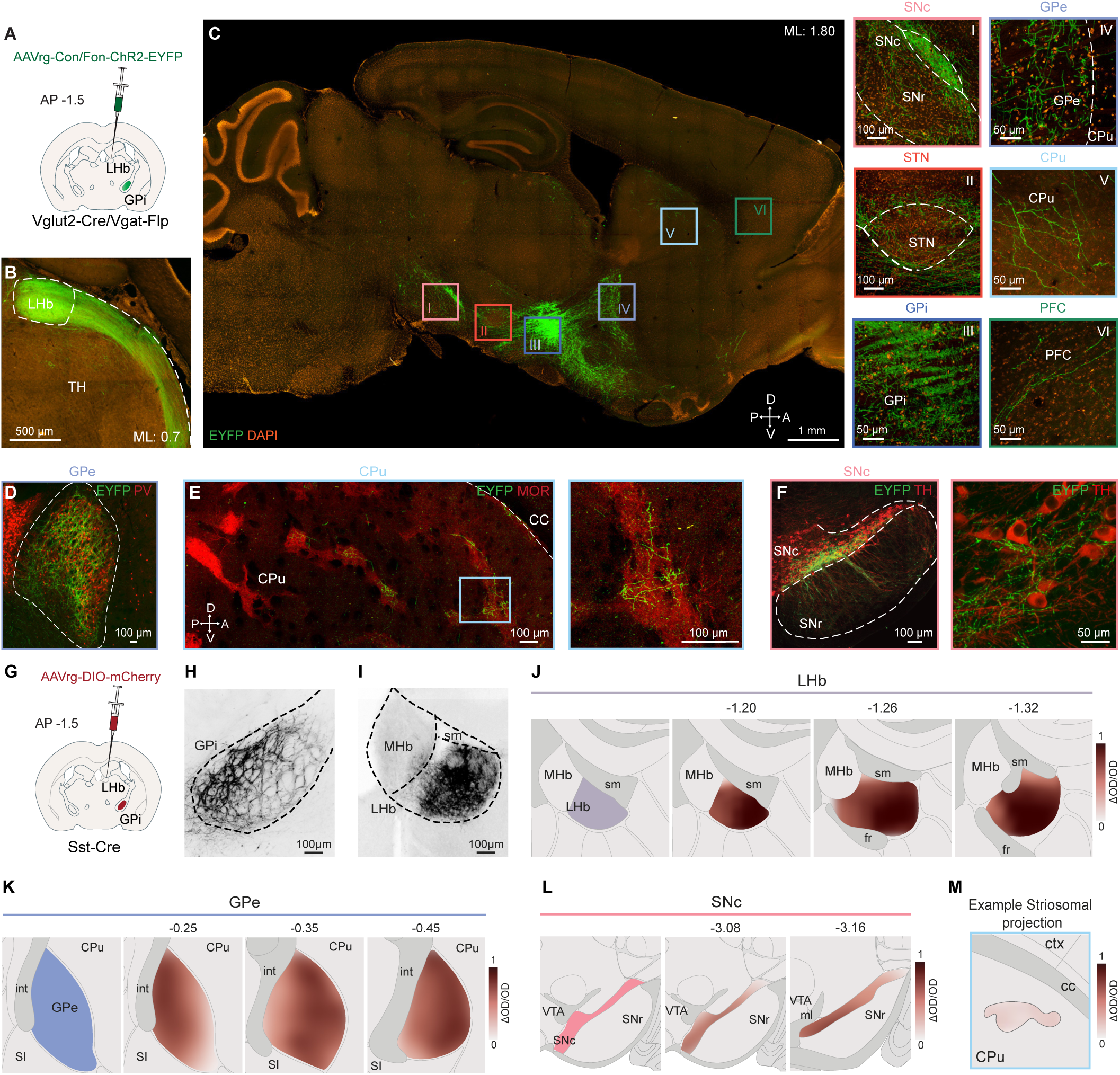
Mapping of Collateral projections of LHb-projecting Vglut2+/Vgat+/Sst+ GPi neurons. **A**, Schematic representation of the experimental strategy (Vglut2/Vgat^GPi-LHb^): AAVrg-Con/Fon-ChR2-EYFP virus was injected in the Lhb of Vglut2-Cre/Vgat-Flp animals. **B**, Sagittal section showing the projections of Vglut2+/Vgat+^GPi-LHb^ neurons in LHb. N=7 animals. **C**, Left: sagittal section showing the collateral projections of Vglut2+/Vgat+^GPi-LHb^ neurons in different brain regions. Roman numerals indicate regions of interest. Right: close-up images of Vglut2+/Vgat+^GPi-LHb^ cell bodies in the GPi (III) and axonal projections in the SNc (I), STN (II), GPe (IV), CPu (V) and PFC (VI). N=7 animals. **D**, ISH showing the Vglut2+/Vgat+^GPi-LHb^ axons in the GPe confirmed by parvalbumin (PV) immunostaining. **E**, Left: ISH demonstrating Vglut2+/Vgat+^GPi-LHb^ axonal collaterals targeting striosomes in the CPu. Striosomes identified by μ-Opioid Receptor 1 (MOR) staining. Right: Close up of Vglut2+/Vgat+^GPi-LHb^ axons projecting in one striosome area. **F**, Left: ISH depicting Vglut2+/Vgat+^GPi-LHb^ axons in the SNc confirmed by tryptophan hydroxylase (TH) staining. Right: Close up of Vglut2+/Vgat+^GPi-LHb^ axons surrounding TH-labelled neurons. **G**, Schematic representation of an alternative experimental strategy (SST^GPi-LHb^) to achieve selective expression in LHb-projecting GPi neurons: injection of AAVrg-DIO-mCherry into the LHb of Sst-Cre mice. **H**, Coronal section showing the labelling of SST^GPi-LHb^ neurons in GPi. N=3 animals. **I**, Coronal section showing the projections of SST^GPi-LHb^ axons in LHb. N=3 animals. **J**, Heatmap showing the average fluorescence intensity of the projections of SST^GPi-LHb^ axons in three anteroposterior levels of LHb. **K**, Heatmap showing the average fluorescence intensity of the projections of SST^GPi-LHb^ axons in three anteroposterior levels of GPe. **L**, Heatmap showing the average fluorescence intensity of the projections of SST^GPi-LHb^ axons in two anteroposterior levels of SNc. **M**, Example heatmap showing the fluorescence intensity of the projections of SST^GPi-LHb^ axons in one striosome of one Sst animal. Abbreviations: AP: anteroposterior, GPi: globus pallidus interna, LHb: Lateral habenula, TH: thalamus, ML: mediolateral, EYFP: enhanced yellow fluorescent protein, SNc: substantia nigra pars compacta, SNr: substantia nigra pars reticulata, GPe: globus pallidus externa, STN: subthalamic nucleus, CPu: Caudate putamen, PFC: prefrontal cortex, MHb: Medial habenula, sm: stria medullaris, fr: fasciculus retroflexus, OD: optical density, int: internal capsule, SI: substantia innominata, VTA: ventral tegmental area, ml: medial lemniscus, ctx: cortex, cc: corpus callosum.

To validate these findings, we next used a second genetic targeting strategy based on the specific expression of Sst in GPi-LHb neurons^13,15^. In this strategy, we used Sst-Cre mice and injected retrograde AAVs into the LHb to achieve Cre-dependent labeling of GPi-LHb neurons based on their Sst expression (experimental strategy referred to as Sst^GPi-LHb^, N = 3 mice; **Figure 1G-I and Supplementary Figure 2K-T**). We found that this strategy confirmed our findings on the extensive collateralized network of GPi-LHb neurons (i.e. their projections to GPe, CPu striosomes, SNc), and also eliminated the unspecific labeling of sparse neurons in SI, VTA, or DR compared to the Vglut2/Vgat^GPi-LHb^ neuron targeting strategy. We further quantified the GPi-LHb axonal intensity in the targeted regions using the Sst^GPi-LHb^ labelling strategy (N = 3), which showed the highest projection density in the LHb, followed by projections density in the GPe, and with lower projection densities in the SNc and in striosomes (**Figure 1J-M** and **Supplementary Figure 3A-D**).

To further confirm our findings on the feedback organization of GPi-LHb projections, we established a third genetic labeling strategy: we injected a Cre- and Flp-dependent viral construct (AAV8-Con/Fon-ChR2-EYFP) directly into the GPi of Vglut2-Cre/Vgat-Flp mice (experimental strategy referred to as Vglut2/Vgat^GPi^; N = 6 mice; **Supplementary Figure 2U**). This direct injection approach again labeled projections to the GPe and striosomes (**Supplementary Figure 2V-AD**). However, we observed some additional axonal labeling in the subthalamic nucleus (STN) and substantia nigra pars reticulata (SNr), which was inconsistent with the labeling profile in the other two strategies (**Supplementary Figure 4A-C**). To determine this projections profile in SNr was the result labeling of Vglut2/Vgat-expressing neurons in the STN, we injected a retrograde Cre- and Flp-dependent viral construct (AAVrg-Con/Fon-ChR2-EYFP) in the SNc of Vglut2-Cre/Vgat-Flp mice. We found that this strategy resulted in the retrograde labeling of cells in the STN, confirming that Vglut2/Vgat+ STN neurons form a direct STN-SNr projection, which is distinct from the GPi-LHb pathway (**Supplementary Figure 4D-F**).

To investigate whether individual GPi-LHb neurons collateralized to multiple basal ganglia target regions, we labeled GPi-LHb neurons to determine their collateral projections to two targets: the SNc and the GPe. In this strategy, we injected mice with three AAVs: a retrograde Cre-expressing virus (AAVrg-Cre) in the LHb, a retrograde Cre-dependent EGFP-expressing virus (AAVrg-DIO-EGFP) in the SNc, and a retrograde Cre-dependent mCherry-expressing virus (AAVrg-DIO-mCherry) in the GPe (N=3; **Supplementary Figure 5A**). Using this triple-virus retrograde labeling strategy, we found single Sst+ GPi-LHb neurons labeled with both EGFP (i.e., projecting to SNc) and mCherry (i.e., projecting to GPe), as well as their EGFP+/mCherry+ axons in LHb (**Supplementary Figure 5A-B**). To further confirm the collateralization of single GPi-LHb neurons, we established another intersectional viral strategy in Sst-Cre mice (N = 3) using injections of three different AAVs: a retrograde AAV with Cre-dependent expression of tetanus toxin (AAVrg-DIO-TetLCh-dTomato) in the LHb, a retrograde AAV with Cre-dependent expression of the Flp recombinase (AAVrg-DIO-FLPO) in the GPe, and a retrograde AAV with Cre- and Flp-dependent EGFP expression (AAVrg-CreOn-FlpOn-EGFP) in the SNc (**Supplementary Figure 5C**). This strategy resulted in expression of EGFP and dTomato in single GPi neurons and labeled axons in both the LHb and GPe, supporting the extensive collaterals of GPi-LHb neurons to the GPe and the SNc (**Supplementary Figure 5D-E**). In conclusion, our results show that GPi-LHb neurons form an extensive and complex feedback network in the basal ganglia, with collateral projections that target key circuit nodes: the GPe, the CPu striosomes, and the SNc.

### GPi-LHb neurons synapse and co-transmit glutamate and GABA in basal ganglia targets

We further examined the postsynaptic targets of GPi-LHb neurons performing a series of monosynaptic retrograde tracing experiments using modified rabies virus and patch-clamp recordings. To assess the postsynaptic specificity of GPi-LHb neurons across distinct striatal populations, investigate whether GPi-LHb neurons preferentially form synapses on striatal projection neurons (SPNs) in striosomes (i.e. Oprm1+) versus D2-expressing SPNs (iSPNs), we injected a helper virus (AAV-DIO-TVA-V5-RbG), followed by a modified rabies virus expressing EGFP into the CPu of Oprm1-Cre (N = 6 mice) or A2A-Cre mice (N = 4 mice; **Figure 2A**). The Oprm1-Cre line labels striosomal SPNs, whereas the A2A-Cre line labels iSPNs distributed across both striosome and matrix compartments, allowing us to determine whether GPi-LHb neurons preferentially target striosomal versus D2-expressing SPNs independent of compartmental localization. We found that Sst+ GPi neurons synapse directly and predominantly onto striosomal Oprm1+ SPNs, but not onto A2A-expressing SPNs, highlighting a clear cell-type and compartment-specific preference (**Figure 2A-C**, **Supplementary Figure 6A-B**, and **Supplementary Table 1**). These results are consistent with the projection pattern observed in striosomes and confirm that GPi-LHb neurons preferentially target D2-negative (putative D1) SPNs. Similarly, we investigated whether GPi-LHb neurons directly target dopamine neurons in the SNc, using the modified rabies virus monosynaptic tracing strategy in DAT-Cre mice (N = 3 mice; **Figure 2D**). We found that dopamine neurons (i.e. DAT+) in SNc received monosynaptic inputs from Sst+ GPi neurons thereby confirming a direct synaptic connection (**Figure 2D-F**, and **Supplementary Table 1**).

**Figure 2.**
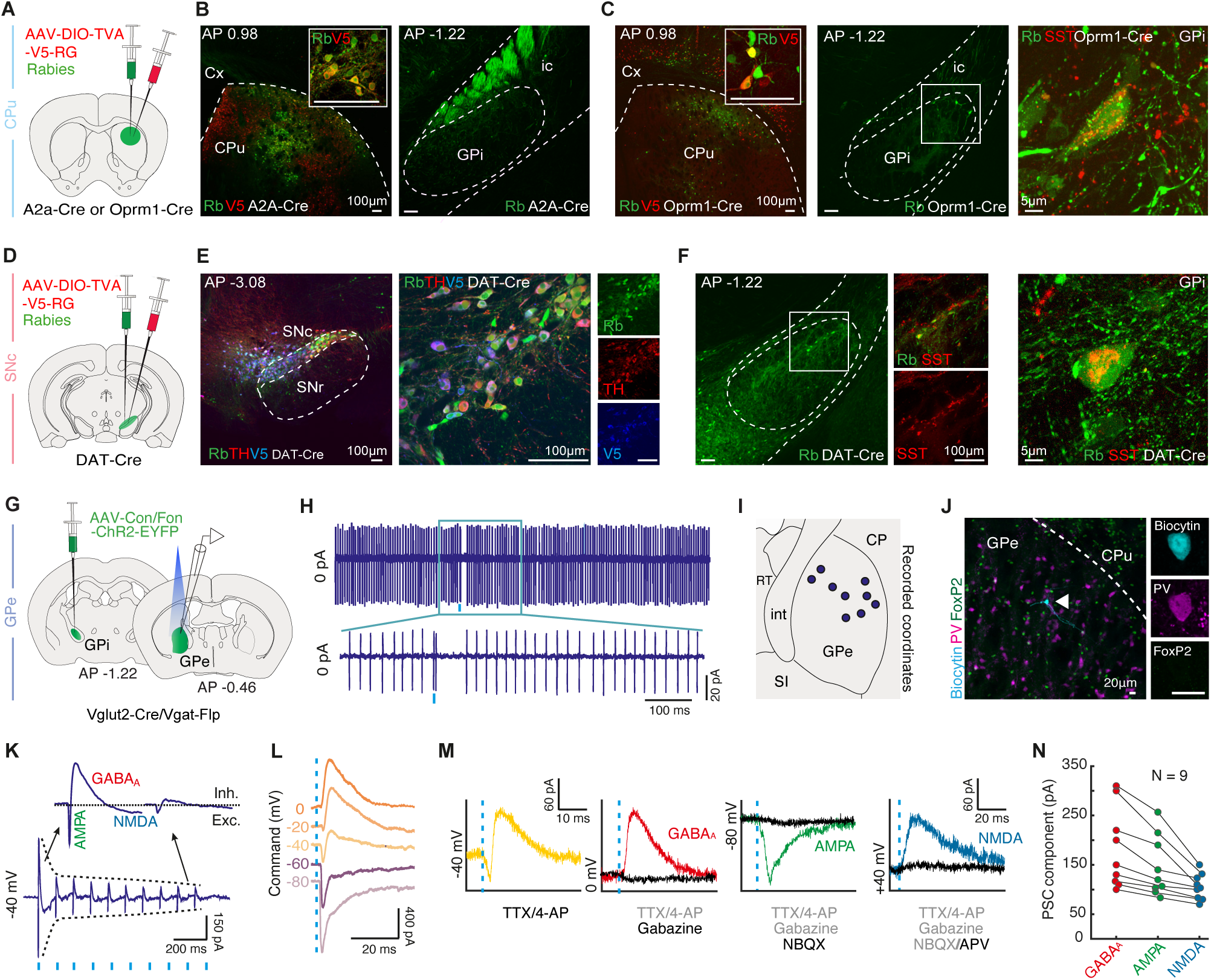
GPi-LHb neurons make monosynaptic connections in the basal ganglia and co-transmit glutamate and GABA. **A**, Schematic illustration of the experimental strategy: AAV-DIO-TVA-V5-RbG (V5) and Rabies-EGFP (Rb) were injected into the dorsolateral CPu of A2a-Cre and Oprm1-Cre mice, 21 days apart. **B**, Representative ISH image of starter neurons co-expressing V5 (red) and Rb (green) in the CPU of an A2a-Cre mouse. Insert: close-up of starter neurons. Right: No Rb+ neurons in the GPi. Scale bars: 100μm. Replicated in N = 4 A2a-Cre mice; for cell counts see **Supplementary Table 1.** See also **Supplementary Figure 6A**. **C**, Representative image of starter neurons co-expressing V5 (red) and Rb (green) in the CPU of an Oprm1-Cre mouse. Insert: closeup of starter neurons. Middle: Rb+ presynaptic neurons in the GPi. Right: Sst+ (red) Rb+ (green) neuron in the GPi. Scale bars: 100μm (left, middle); 5μm (right). Replicated in N = 6 Oprm1-Cre mice; for cell counts see **Supplementary Table 1**. See also **Supplementary Figure 6B**. **D**, Schematic illustration of the experimental strategy: AAV-DIO-TVA-V5-RbG (V5) and Rabies-EGFP (Rb) were injected into the SNc of DAT-Cre mice, 21 days apart. **E**, Representative ISH image of starter neurons co-expressing the V5 (blue), Rb (green), and TH (red) in the SNc of a DAT-Cre mouse. Right: closeup of starter neurons. Scale bars: 100μm. **F**, Representative Rb+ GPi neurons (green) targeting DAT+ SNc neurons. Close-up images to the right show Sst+ (red) Rb+ (green) GPi neurons. Right: Close-up of one Sst+ Rb+ GPi neuron. Scale bars: 100μm (left), 5μm (right). Replicated in N = 3 DAT-Cre mice; for cell counts see **Supplementary Table 1**. **G**, Schematic illustration of the experimental strategy: AAV-Con/Fon-ChR2-EYFP was unilaterally injected directly into the GPi of Vglut2-Cre/Vgat-Flp mice. ChR2-expressing GPi axon terminals were activated by blue light and postsynaptic currents in GPe neurons were recorded in whole-cell voltage-clamp mode. **H**, Optogenetic activation (3ms, blue tick mark) of GPi axons results in an action potential doublet followed by a silent phase in a postsynaptic GPe neuron. Replicated in n = 3 neurons. **I**, Schematic illustration showing the location of each recorded GPe neuron responsive to GPi axonal activation. n = 9 neurons. **J**, ISH image showing a responsive GPe neuron (white arrowhead) filled with biocytin (cyan), co-expressing PV (pink) but not FoxP2 (green). Right: Close-up ISH images of the responsive PV+ FoxP2- GPe neuron. Scale bars: 20μm. Replicated in n = 9 recorded responsive GPe neurons. See also **Supplementary Figure 6D**. **K**, Graph showing the response of a GPe neuron to optogenetic stimulation (10 Hz, 3 ms, blue tick marks) of GPi axon terminals. The envelope (dashed line) indicates a symmetrical short-term depression of the GPi-GPe synapse. Upper traces: Excitatory (Exc, AMPA, NMDA) and inhibitory (Inh, GABAA) postsynaptic current components of the first and tenth pulse. Replicated in n = 3 neurons. Holding potential was -40 mV. See also **Supplementary Figure 6C**. **L**, Graph depicting the optogenetically evoked postsynaptic currents at different holding potentials. Averages of 5 traces for n = 9 neurons. **M**, Graphs depicting the pharmacological dissection of postsynaptic currents evoked by opto-stimulation (blue dashed line). Gabazine (10 μM), NBQX (20 μM) and APV (50 μM) were applied sequentially. TTX (1 μM) and 4-AP (5 mM) were applied throughout the entire recording. Averages of 5 traces for n = 9 neurons. **N**, Graph showing the maximal current amplitude of GABAA, AMPA- and NMDA-currents measured at the holding potentials shown in **M**. Abbreviations: CPu: Caudate putamen, Cx: Cortex, GPi: Globus pallidus interna, ic: internal capsule, SNc: substantia nigra pars compacta, SNr: substantia nigra pars reticulata, GPe: Globus pallidus externa, RT: Reticular nucleus of the thalamus, int: internal capsule, SI: Substantia innominata, CP: Caudate putamen, FoxP2: Forkhead box P2.

GPi-LHb neurons have been shown to co-transmit GABA and glutamate in the LHb^14^. To establish the GABA/glutamate synaptic co-transmission profile of GPi-LHb neurons in other target regions, we investigated the synaptic transmission of Vglut2/Vgat^GPi-LHb^ axons terminals in the GPe. We used optogenetic activation of Vglut2/Vgat^GPi^ axon terminals with electrophysiology recordings from GPe neurons. A Cre- and Flp-dependent viral construct expressing ChR2 (AAV8-Con/Fon-ChR2-EYFP) was injected directly into the GPi of Vglut2-Cre/Vgat-Flp mice, and GPe neurons were recorded using cell-attached and whole-cell patch clamp recordings (**Figure 2G**). The electrophysiological response to optogenetic Vglut2/Vgat^GPi^ axon activation, location, and molecular profile of each patched GPe neuron was characterized (**Figure 2H-J)**. In cell-attached recordings, the optogenetic activation of Vglut2/Vgat^GPi^ axons (3 ms, blue light) resulted in an action potential doublet followed by firing pause, which interrupted fast tonic firing of GPe neurons (**Figure 2H**). This indicates the presence of both excitation and inhibition in an intact GPe neuron. The whole-cell recorded postsynaptic response of GPe neurons to optogenetic activation was monosynaptic (i.e., in TTX, 4-AP) with a delay of 2-3 ms. The response exhibited both excitatory and inhibitory components, and repeated optogenetic stimulation at 10 Hz induced short-term synaptic depression in both (**Figure 2K-L**). Notably, the strong correlation in the jitter of GABA and AMPA currents suggests a high likelihood of co-release, indicating that excitatory and inhibitory neurotransmission originate from the same presynaptic terminals (**Supplementary Figure 6C**). Sequential pharmacological dissection (Gabazine 10uM, NBQX 20uM, AP-V 50uM) confirmed the synaptic co-transmission of glutamate and GABA from Vglut2/Vgat^GPi^ terminals in all GPe neurons with optogenetically evoked postsynaptic response (N = 9 neurons; **Figure 2M-N**). We further probed the subtype identity of the responsive GPe neurons, and we found that all neurons were Pv+, lacked Foxp2 expression (**Figure 2J** and **Supplementary Figure 6D**), and displayed intrinsic electrophysiological properties characteristic of prototypic PV+ GPe neurons^20^ (**Supplementary Figures 6E-F**). These recordings establish the synaptic connection and co-transmission of GABA and glutamate between Vglut2+/Vgat+ GPi neurons and GPe neurons.

To investigate the glutamate/GABA profile of Vglut2+/Vgat+ GPi neurons projections to other basal ganglia targets, we examined the expression of the vesicular transporters for glutamate (i.e., Vglut2) and GABA (i.e., Vgat) in the synaptic boutons of genetically labeled Vglut2/Vgat^GPi-LHb^ axons. To visualize the boutons of Vglut2/Vgat^GPi^ axons, we injected a Cre- and Flp-dependent AAV virus (AAV8-Con/Fon-GCaMP6m) into the GPi of Vglut2-Cre/Vgat-Flp mice and mapped the localization of Vglut2 and Vgat using immunostaining (**Supplementary Figure 6G-H**). In all target regions, most GFP-labeled boutons from Vglut2/Vgat^GPi^ axons contained both Vglut2 and Vgat (co-labeled boutons/total boutons: LHb 433/465; GPe 274/292; CPu 67/75; PFC 31/34; **Supplementary Figure 6I**), supporting that Vglut2/Vgat^GPi^ axons can co-release glutamate and GABA in all target regions. In addition, to estimate the physiological influence of GPi-LHb transmission we quantified the density of Vglut2 and Vgat-expressing boutons in each target region. The most boutons per μm^2^ were found in the LHb, followed by the GPe, CP and lastly the PFC (**Supplementary Figure 6I**). These results further demonstrate that GPi-LHb neurons have the potential to modulate the activity of prominent basal ganglia regions by local co-transmission of glutamate and GABA.

### Sst+ GPi-LHb activity is not essential for movement execution

Based on the projection profile of GPi-LHb neurons targeting key basal ganglia nodes, we first aimed to define their role in spontaneous motor behaviors. We therefore recorded the activity of Sst+ GPi-LHb neurons during exploration in an open field arena using fiber photometry and the genetically encoded calcium indicator GCaMP7s^21^ (**Figures 3A-B**). Sst+ GPi-LHb neurons were labeled by injecting a retrograde AAV driving Cre-dependent expression of jGCaMP7s (AAVrg-DIO-jGCaMP7s) into the LHb of Sst-Cre mice (N = 5, **Figure 3A** and **Supplementary Figure 7A**), as this cell-type specific strategy resulted in the most precise targeting of the GPi-LHb population (**Supplementary Figure 1-4**). The classically motor-associated GPi is composed of Pv+ neurons that project to the thalamus and brainstem^15^, and we therefore used the activity of Pv+ GPi neurons for comparison in our assessment of movement-related activity of Sst+ GPi-LHb neurons. Pv+ GPi neurons were targeted using cell type-specific labeling following injections of an AAV with Cre-dependent jGCaMP7s expression (AAV-DIO-jGCaMP7s) into the GPi of Pv-Cre mice (N = 3, **Figure 3A** and **Supplementary Figure 7B**). Mouse behavior was recorded with a bottom-up camera and pose estimation of three body parts (snout, gravity center, and tail base) was performed using DeepLabCut^22^ (DLC) to identify basic locomotor-related activity in the two discrete neuronal populations (**Figure 3B**).

**Figure 3.**
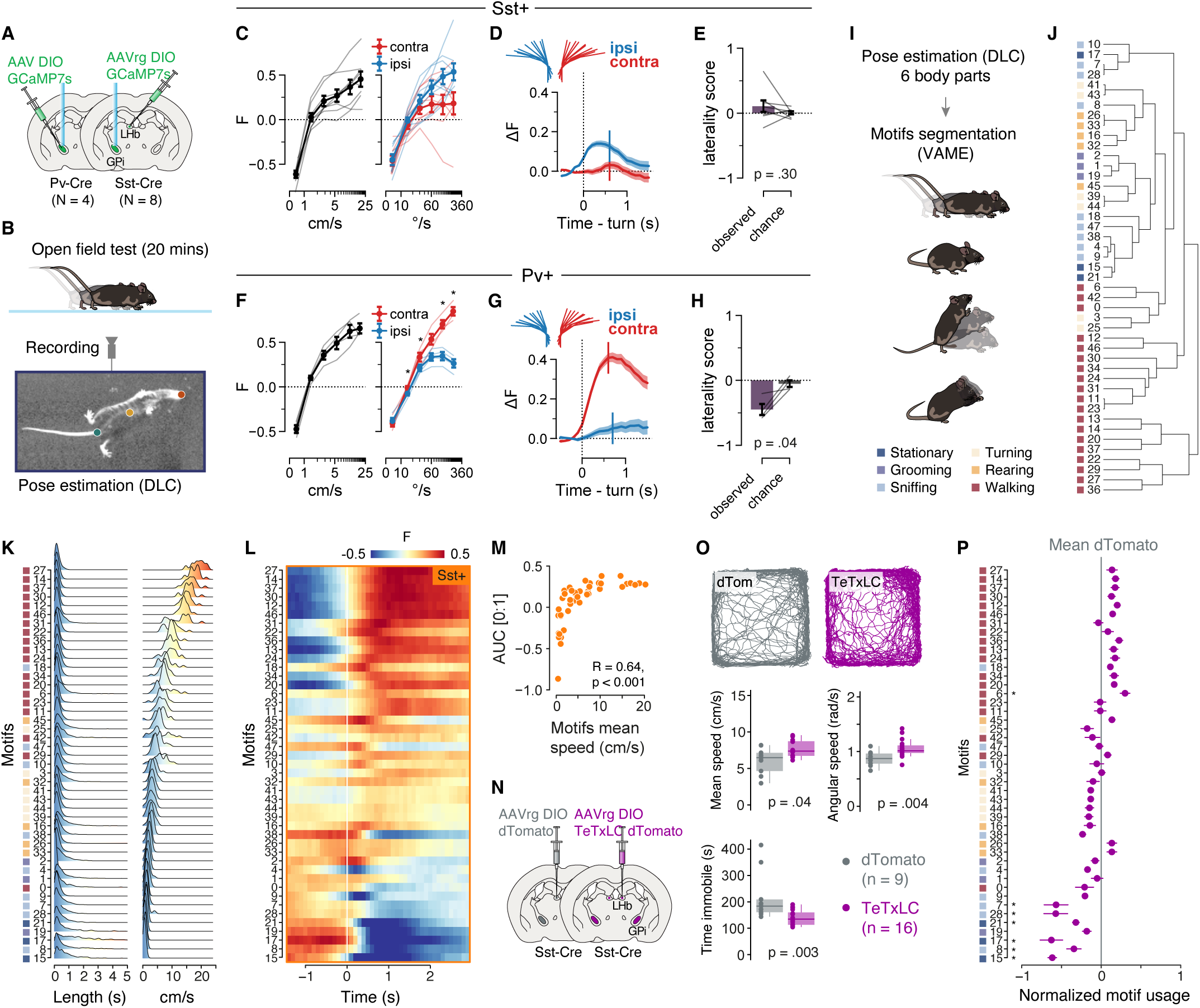
Sst+ GPi neuronal activity correlates with action speed. **A**, Schematic illustration showing the experimental strategy: AAVrg-DIO-jGCaMP7s was injected unilaterally in the LHb of Sst-Cre mice (N = 8). AAV-DIO-jGCaMP7s was injected unilaterally in the GPi of Pv-Cre mice (N = 4). 200μm fibers were implanted above the GPi. See also **Supplementary Figure 7A-B** and **Supplementary Table 2**. **B**, Schematic representation of the open field test setup with a bottom-up camera (top), example tracking of mouse behavior for video analysis and pose estimation of three body parts through DeepLabCut (bottom). **C**, Population activity (F) at different velocities (left) and angular velocities split by turn direction (right) in the open field. Bins spaced evenly along the log-scaled x-axes. Thin lines: individual Sst-Cre mice. Circles: mean ± SEM. Contra- vs ipsiversive turn F: all bins p > 0.3; paired t-tests, Holm-Sidak correction. N = 8 mice. **D**, ΔF during spontaneous ipsi- and contraversive turns in the open field, aligned to turn onset. Baseline for ΔF: mean F of the 0.5s preceding the onset of the turn. Traces: mean ± SEM. Vertical lines: median turn offset times. Ipsi/contra n = 1235/1360 turns (pooled, N = 8 mice). **E**, Measure indicating how well single-turn mean ΔF discriminates turn direction. The laterality score was defined as the area under the ROC curve (AUC), scaled from 1 to 1. Positive vs negative scores indicate relatively higher activation during ipsi- vs contraversive turns. Chance level established by shuffling the turn direction labels prior to score computation. Gray lines: individual Sst-Cre mice. Bars: mean ± SEM. p = 0.30; paired t-test. N = 8 mice. **F-H**, same as C-E for Pv-Cre mice. **F**, *: p < 0.05; paired t-tests, Holm-Sidak correction. N = 4 mice. **G**, Ipsi/contra n = 355/395 turns (pooled, N = 4 mice). **H**, p = 0.04; paired t-test. N = 4 mice. **I**, Schematic illustration of the video analysis: pose estimation of six body parts and segmentation of behavior motifs. **J**, Dendrogram reporting the clustering and behavioral annotations of the 45 identified motifs. **K**, Densities plots reporting the motifs length (left) and the mouse speed (right) during the execution of the motifs. **L**, Mean average of calcium transients signals of Sst+ GPi-LHb population recorded through fiber photometry and aligned to motifs onset. The motifs were ordered according to the motif’s speed (cf. panel **J**). **M**, AUC of the signal (period between 0 and +1 sec) correlated with the motif’s speed for Sst+ GPi-LHb recording. Spearman’s correlation coefficients are reported. **N**, Illustration showing the experimental strategy: AAVrg-DIO-dTomato (N = 9) or AAVrg-DIO- TeTxLC-dTomato (N = 16) were injected bilaterally in the LHb of Sst-Cre mice (see also **Supplementary Figure 7C**). **O**, Upper panel: example traces of a control and TeTxLC mouse. Bottom panels: boxplots reporting linear and angular speed, and the time passed immobile for both control and TeTxLC mice. The p-values of the Mann-Whitney U test are reported in each panel. **P**, Normalized (on the performance of control mice) motifs usage for TeTxLC mice. Multiple Mann-Whitney U test followed by Holm-Sidak correction. Mean ± SEM. *: p<0.05. See **Supplementary Table 3** for more information about statistical tests. Abbreviations: ΔF: delta fluorescence, ROC: Receiver operating characteristic, AUC: area under curve.

We first observed that Sst+ GPi population activity increased monotonically with movement velocity, similar to the Pv+ GPi population activity (**Figure 3C** and **F**). However, while the Pv+ GPi population activity showed a strong and consistent preference for contraversive over ipsiversive turns (relative to the recorded hemisphere), no clear preference could be established for the Sst+ GPi population (**Figure 3C-H**). The motor-related signals of Sst+ GPi were therefore discretely different from those of the canonical motor GPi output, suggesting that the two GPi outputs may shape actions and motor programs through distinct mechanisms.

We therefore decided to investigate the activity of Sst+ GPi-LHb neurons in more detail during specific actions and motor motifs. We trained a second DLC model with six markers (four paws, snout, and tail base) and the behavioral repertoire was categorized into discrete motifs through unsupervised learning using a variational autoencoding motif extractor^23^ (VAME, **Figure 3I**). This classification identified 45 unique motor motifs during open field exploration. Motifs typically lasted less than a second and were ranked by their average speed (**Figure 3J-K**). When we aligned the calcium signals from Sst+ GPi-LHb neurons with the onset of each motif, we found that low-speed motifs showed a strong decrease in neuronal activity, whereas the high-speed motifs showed an increase in activity, independently of the type of motor action performed (**Figures 3L-M**). In summary, recording of Sst+ GPi neurons during self-paced exploration revealed a preferential activation during vigorous and high-speed actions.

To directly test the causal role of Sst+ GPi-LHb neurons in the execution of high-speed motor motifs, we used a neurotransmitter release inactivation strategy based on cell type-specific viral expression of the tetanus toxin light chain^24^ (TeTxLC). This method was chosen for its ability to achieve robust and sustained suppression of synaptic transmission in targeted neuronal populations, making it particularly suitable for long-term behavioral studies (see **Supplementary information** for more details). Specifically, we injected a retrograde AAV with Cre-dependent expression of TeTxLC (AAVrg-DIO-TeTxLC-dTomato) bilaterally into the LHb of Sst-Cre mice (N = 16 mice, **Figure 3N** and **Supplementary Figure 7C**). This experimental group was compared with a control group injected with a retrograde AAV with dTomato expression (N = 9, AAVrg-DIO-dTomato). Unexpectedly, we found that genetic inactivation of the Sst+ GPi-LHb neurons resulted only in a small increase in the linear and angular speed as well as a small reduction in immobility (**Figure 4O**). We then analyzed the behavior based on the unsupervised classification and found a significant reduction in the usage of low-speed motifs following the inactivation of Sst+ GPi-LHb neurons (**Figure 4P**). Overall, this inactivation experiment suggests that although Sst+ GPi-LHb neurons are specifically activated during high-speed motor motifs, their activity is not necessary for initiating movements and does not directly control the execution of vigorous motor actions.

**Figure 4.**
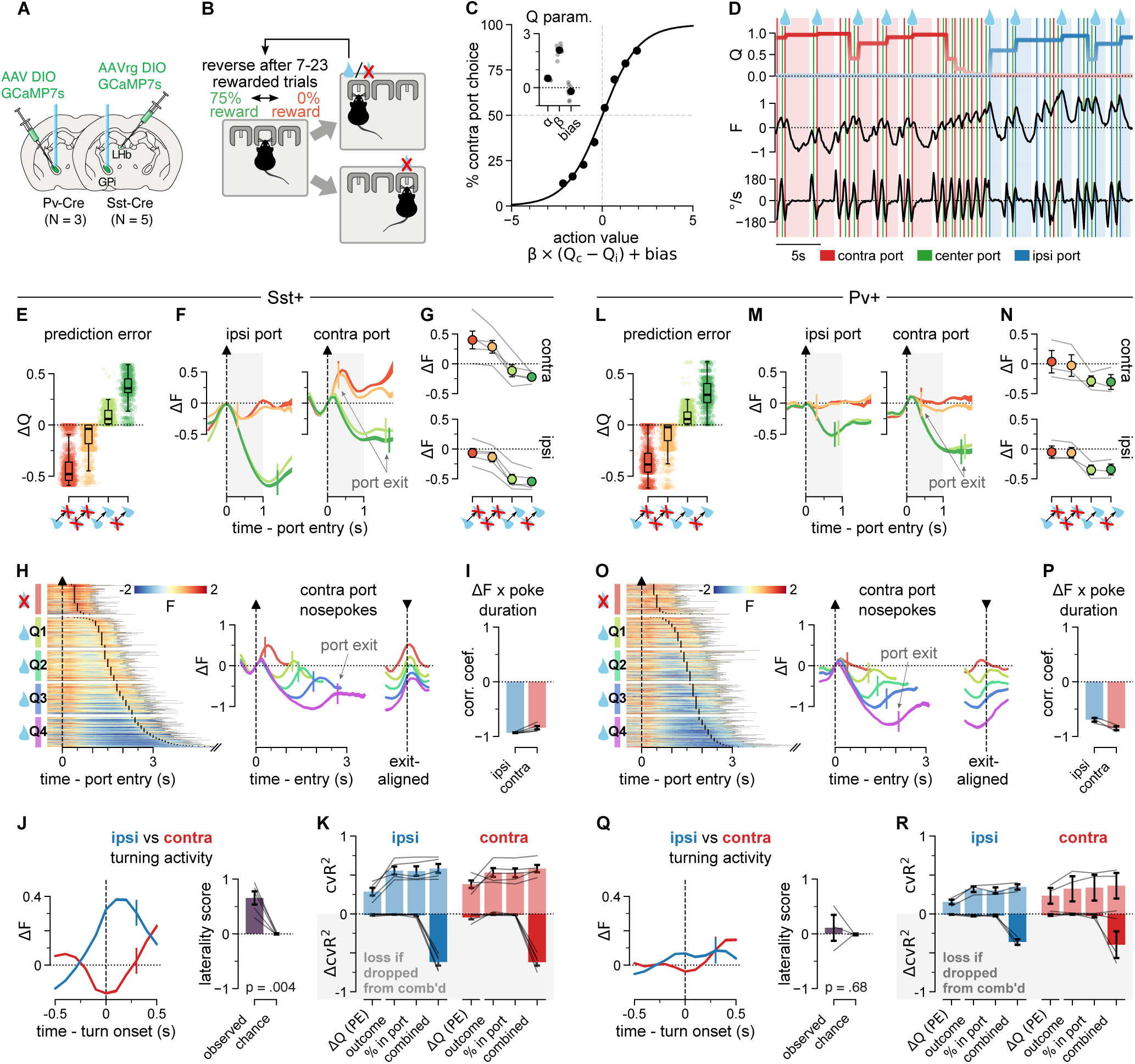
Sst+ GPi-LHb population activity is movement-related and ipsiversive turn-preferring. **A**, Top: Schematic representation of the experimental strategy: AAVrg-DIO-jGCaMP7s was injected unilaterally in the LHb of Sst-Cre mice (N = 5). AAV-DIO-jGCaMP7s was injected unilaterally in the GPi of Pv-Cre mice (N = 3). 200μm optic fibers (OF) were implanted above the GPi. Bottom: jGCaMP7s (green) expressed in the GPi of a Sst-Cre mouse. See also **Supplementary Figure 7A-B** and **Supplementary Table 2**. **B**, Probabilistic reversal task. After trial initiation at the center nose-poke port, a reward becomes available (with 75% probability) at one of the two peripheral ports. The rewarded choice reverses unpredictably every 7-23 rewarded trials. **C**, Graph showing the probability of contraversive port choice (relative to implanted hemisphere) as a function of the outcome history-based action value. Action values were fitter per mouse using Q-learning. Line: logistic regression fit, trials pooled over mice. n = 27053 trials (pooled, N = 8 mice; 3 Pv-Cre + 5 Sst-Cre). Circles: mean ± SEM of mice for 8 evenly sized action value bins. N = 8 mice. Inset: Q-learning parameters. Gray circles: individual mice. Black circles: mean ± SEM. N = 8 mice. **D**, Exemplary session snippet showing Q value estimates for both port choices, Sst+ GPi-LHb population activity, and the mouse’s angular velocity over the course of several trials. Lines: nose-poke port entries. Light shading: port occupation. All color coding as indicated in the panel legend. The water drop icon marks rewarded choice port entries; unmarked entries were not rewarded. Data from one Sst-Cre mouse. **E**, Graph depicting trial-by-trial changes in the fitted port choice Q values (ΔQ, i.e. prediction error). Trials split by current and previous trial outcomes. Open circles: single trials. Boxplots: center lines represent the distribution median; edges, the upper and lower quartiles; whiskers, the minimum and maximum values within the quartiles ± 1.5 x IQR. n = 1220/1867/1157/606 trials (pooled, N = 5 mice). **F**, Graph showing the fluorescence change (ΔF) following contraversive (left) and ipsiversive (right) choice port entries (time 0, dashed arrows). Trials split by current and previous trial outcomes. Baseline for ΔF: F at port entry. Color coding as in panel E. Traces: mean ± SEM. Vertical lines: median port exit times. Gray shading: 1s time window used to compute means in panel G. Ipsi n = 613/983/498/296, contra n = 607/884/659/310 trials (pooled, N = 5 mice). **G**, Graph demonstrating the mean ΔF in the second after choice port entry (cf. panel F). Trials split by the current and previous trial outcome. Gray lines: individual Sst-Cre mice. Circles: mean ± SEM. A repeated-measures ANOVA showed significant main effects for the current trial outcome (p < 0.001) and the port side (p = 0.004). The main effect of the previous trial outcome (p = 0.097) and all interaction effects were insignificant (all p > 0.3). N = 5 mice. **H,** Raster: single-trial fluorescence (F) time-locked to contraversive choice port entry. Rewarded trials were split into nose-poke duration-based quartile bins (Q1-4) and a fifth, equivalently sized bin of unrewarded trials was randomly sampled. All bins sorted by poke duration. Black ticks mark port exit times. Traces: ΔF means ± SEM for each trial bin of the raster plot, plotted aligned to port entry (left, dashed up-arrow) and port exit times (right, dashed down-arrow). Baseline for ΔF: F at port entry. Vertical lines: median port exit times. n = 242 trials per bin (pooled, N = 5 animals). **I**, Pearson correlation between mean ΔF and time spent nose-poking for all ipsiversive and contraversive choice port nose-pokes, irrespective of trial outcome. Gray lines: individual Sst-Cre mice. Bars: mean ± SEM. N = 5 mice. **J**, Left: ΔF during task-engaged ipsi- and contraversive turns, aligned to turn onset. Includes choice and initiation turns. Baseline for ΔF: mean F of the 0.5s preceding the onset of the turn. Traces: mean ± SEM. Vertical lines: median turn offset times. Ipsi/contra n = 4658/4612 turns (pooled, N = 5 mice). Right: ROC-AUC score-based laterality score indicating how well single-turn mean ΔF discriminates turn direction. Positive vs negative scores indicate higher activation during ipsi- vs contraversive turns. Chance level established by shuffling the turn direction labels prior to score computation. Gray lines: individual Sst-Cre mice. Bars: mean ± SEM. p = 0.004; paired t-test. N = 5. **K**, Predictive power of ridge regressions predicting trial-by-trial mean ΔF in the 1s window following ipsi or contra choice port entry (see panel F), fit per mouse and turn direction. cvR^2^: cross-validated variance explained by each variable individually (ΔQ, outcome, % in port) and combined. ΔcvR^2^: loss of explanatory power of the combined 3-variable model when variables are individually and jointly scrambled. Gray lines: individual Sst-Cre mice. Bars: mean ± SEM. Pairwise comparisons of the single-variable models’ cvR^2^ scores were insignificant (all p > 0.3), except for the ipsi port comparisons involving the “ΔQ” model (both p < 0.03; paired t-tests, Holm-Sidak correction). Exclusion of individual variables from the combined model failed to affect its performance (all p > 0.07; t-tests, Holm-Sidak correction). N = 5 mice. **L- R**, same as E-K for Pv-Cre mice. **L**, n = 986/1509/1174/534 trials (pooled, N = 3 mice). **M**, ipsi n = 543/1054/626/249, contra n = 443/455/548/285 trials (pooled, N = 3 mice). **N**, Repeated-measures ANOVA: significant main effect for the current trial outcome (p = 0. 029). The main effect of the previous outcome (p = 0.191) and all other main and interaction effects were insignificant (all p > 0.3). N = 3 mice. **O**, n = 208 trials per bin (pooled, N = 3 mice). **P**, N = 3 mice. **Q**, Left: Ipsi/contra n = 4099/4250 turns (pooled, N = 3 mice). Right: p = 0.68; paired t-test. N = 3. **R**, Pairwise comparisons of the single-variable models’ cvR^2^ scores were insignificant (all p > 0.05; paired t-tests, Holm-Sidak correction). Exclusion of individual variables from the combined model failed to affect its performance (all p > 0.1; t-tests, Holm-Sidak correction). N = 5 mice. See **Supplementary Table 3** for more information about statistical tests. Abbreviations: F: fluorescence, Q: trial-by-trial value estimate of the Q-learning model, PE: prediction error.

### Sst+ GPi-LHb activity does not code prediction errors in a serial reversal task

Since we found that the activity of Sst+ GPi-LHb neurons was positively correlated with vigorous movements but not necessary for their execution, we aimed to further investigate their involvement in action selection and reinforcement learning. Specifically, we sought to investigate whether the Sst+ GPi-LHb population conveys an “anti-reward” outcome evaluation signal—such as negative valence or inverse reward prediction error—as previously proposed^12,16^. To directly test this hypothesis, we recorded the activity of Sst+ GPi-LHb and Pv+ GPi neurons during a two-choice probabilistic serial reversal task using fiber photometry (**Figure 4A**).

In the probabilistic reversal task, water restricted mice faced a choice between two nose-poke ports which flanked a central initiation port (**Figure 4B**). On any given trial, only one of the choice ports yielded a water reward with 75% probability. The location of the reward reversed unpredictably every 7-23 rewarded trials, forcing mice to evaluate trial outcomes in the light of recent history in an ongoing fashion to infer the correct choice^13,16,25,26^. The mice performed this task successfully, allowing us to fit a Q-learning model that accurately captured mouse behavior and generated trial-by-trial value (Q) as well as value update (ΔQ, i.e., prediction error) estimates (**Figure 4C-D**).

To test whether Sst+ GPi-LHb neurons encode negative valence or inverse ΔQ, we split both positive valence (rewarded) and negative valence (non-rewarded) outcome trials into respective low and high ΔQ trials. In low ΔQ trials, the current outcome matched the outcome obtained on the previous trial, in high ΔQ trials it did not. Given the blocked task structure, the current outcome is the best predictor of future outcomes, and matching outcomes are therefore expected to result in minor value updates, whereas mismatched outcomes require reevaluation of the choice–value mapping. This pattern of reevaluation was reflected in the behavior-fit ΔQ estimates, showing that both GPi-recorded mouse groups updated their choice preferences accordingly (**Figure 4E** and **L**).

To define how neuronal activity relates to the inferred value updates, we aligned Sst+ GPi- LHb calcium signals to entry into the choice port (i.e., where the outcome was presented). We found that ipsiversive port entries (“ipsi port”, i.e., ipsilateral relative to the recorded hemisphere) that were rewarded showed a decrease in population activity, while trials with non-rewarded ipsiversive choices showed no marked signal modulation (**Figure 4F-G**). Notably, responses were independent of ΔQ, as both low and high ΔQ trial outcomes showed signals of similar magnitude during the choice port entry (**Figure 4E-G**). Rewarded contraversive port entries (“contra port”) were likewise followed by a signal decrease (albeit weaker than at the ipsiversive port), while non-rewarded contraversive entries were followed by a moderate increase in population activity (**Figure 4F-G**). Like ipsiversive port responses, contraversive port responses were also unaffected by ΔQ (**Figure 4E-G**). To thoroughly evaluate these observations statistically, we conducted a repeated-measures ANOVA examining the effects of the previous and current trial outcomes, and of port choice, on the activity changes (ΔF) recorded in the second following choice port entries (data in **Figure 4G**). We found a significant main effect of the current trial outcome (F_(1,4)_ = 228.13, p = .0001), indicating that Sst+ activity was indeed relatively down-modulated following reward versus non-reward outcomes. Additionally, there was a significant main effect of the port choice (F_(1,4)_ = 35.19, p = 0.004), indicating that Sst+ activity levels were relatively up-modulated in the second after mice entered the contra-versus the ipsiversive choice port. Finally, there was no significant main effect of the previous trial outcome (F_(1,4)_ = 4.65, p = 0.1), nor were there any significant interaction effects among the factors (all p > 0.3). The fact that the previous trial outcome did not significantly modulate Sst+ activity supports that Sst+ GPi activity is not shaped by expectation and therefore does not signal ΔQ.

We hypothesized that the Sst+ distinctive activity peak detected after non-rewarded contraversive choice trials – and indeed the main effects of current outcome and port choice generally – could be motor-rather than valence-related, given our prior finding of a strong correlation between neuronal activity and vigorous movements in the open field. A strong negative correlation between the activity changes captured after choice port entry and the duration of the nose poke (**Figure 4I**) further suggested a link between movement (or immobility) and activity in the reversal task. To directly test this, we analyzed single-trial, outcome-locked fluorescence traces independent of outcome valence. We found that Sst+ GPi-LHb population activity consistently increased whenever mice exited the contra port, irrespective of whether the trial outcome was positive or negative (**Figure 4H**). This supports that the Sst+ GPi activity is associated with the action sequence following the entry in the contra port instead of signaling the negative (i.e., unrewarded) condition. We therefore hypothesized that Sst+ GPi-LHb population activity increased specifically during ipsiversive turns. To test this, we examined how population activity correlated with task-engaged ipsiversive versus contraversive turning. Unexpectedly, we found that the Sst+ GPi-LHb population activity showed marked differences depending on turn direction. Specifically, the activity of Sst+ GPi neurons increased appreciably preceding ipsiversive turns and moderately decreased preceding contraversive turns (**Figure 4J**). This robust hemispheric lateralization of the turn signals was qualitatively different from our observations in the open field, where the Sst+ GPi population was not significantly activated during ipsiversive turns (**Figures 3C-E**).

A comparison of the population activity of Pv+ GPi and Sst+ GPi-LHb neurons in response to choice port entries and exits, revealed qualitatively similar activity following rewarded versus non-rewarded trials. Similar to the Sst+ GPi population, the Pv+ GPi population showed decreased activity following rewarded port entries, with little evidence supporting inverse ΔQ or prediction error coding (**Figure 4L-N**). Accordingly, ANOVA analysis of the Pv+ population activity revealed a significant main effect of the current (F_(1,2)_ = 33.24, p = 0.03), but not of the previous trial outcome (F_(1,2)_ = 3.80, p = 0.19), and no significant interactions between the factors (all p > 0.3; **Figure 4N**). However, unlike Sst+ GPi-LHb neurons, Pv+ GPi neurons showed moderate increase after non-rewarded entries (**Figure 4M**) or exits from the contra or ipsi port (**Figure 4O**), nor did the ANOVA analysis evidence a significant main effect of port choice (F_(1,2)_ = 1.00, p = 0.4). Correspondingly, Pv+ GPi activity decreased during nose poke-related immobility (**Figure 4P**) but lacked significant direction-selective modulations during turns in the serial reversal task (**Figure 4Q**).

Our findings suggest that Sst+ GPi-LHb neurons are specifically activated during task-relevant directional movements, without significant modulation by value and value-updating signals. To directly test whether prediction error (ΔQ) or outcome valence signals can explain any activity modulation in the Sst+ or Pv+ GPi population following choice port-entry, and which is not instead predicted by immobility during nose poking (i.e. the fraction of the 1s averaging window spent nose-poking), we applied a series of ridge regression models (**Figures 4K** and **R**). Across both GPi pathways and port choices, “ΔQ”, “outcome”, and “% in port” single-variable models performed roughly equivalently, with the “ΔQ” model tending to perform slightly worse. Critically, the proportion of the variation explained (*R*^2^) by the better-performing “outcome” and “% in port” models did not differ significantly (all p > 0.6; multiple paired t-tests). Moreover, the “ΔQ”, “outcome” and “% in port” variables proved interchangeable: scrambling any single variable in models which combined all three had minimal impact on the explanatory power of the combined models (all p > 0.07; multiple 1-sample t-tests with Holm-Sidak correction). Supporting the relevance of immobility-related signals in the GPi pathways, we found that the activity of both GPi populations was inversely correlated with immobility during fear conditioning, and that the aversive shock-predicting auditory cues did not modulate either pathway, also supporting the absence of clear valence signals (**Supplementary Figure 8A**).

Overall, we conclude that the activity of the Sst+ GPi-LHb pathways in the serial reversal task was modulated during specific motor actions rather than by the outcome value or prediction errors, and that this activity was strongly lateralized specifically in goal-directed and task-engaged directional movements.

### Adaptation of Sst+ GPi-LHb activity after reward location changes in a single-reversal maze task

Based on our findings, we hypothesized that Sst+ GPi-LHb neurons selectively engage to promote movement or movement-related decisions in a context or task-dependent manner. We therefore aimed to investigate how Sst+ GPi activity developed during the development of goal-directed strategies. In the serial reversal task, mice could have established a switching strategy prior to the start of neural recording, and the fast and constrained movements in the operant chamber could result in overlap of the slower signals from the genetically encoded calcium sensors. To facilitate the identification of behaviorally relevant calcium signals during relevant actions and over learning, we decided to record Sst+ GPi-LHb and Pv+ GPi population activity in a single-reversal task in a larger maze to extend the duration of movement and decision-making signals (**Figure 5A-B**). In this task, water-restricted mice run through a center corridor and make sharp turns into one of the two choice corridors. Mice received rewards when they chose the correct corridor and when they initiated a new trial at the start location. In the first part of this task, water reward was available in the choice corridor requiring contraversive turning relative to the recorded hemisphere, followed by a switch of the reward location requiring ipsiversive turning on the 15^th^ session (**Figure 5C**).

**Figure 5.**
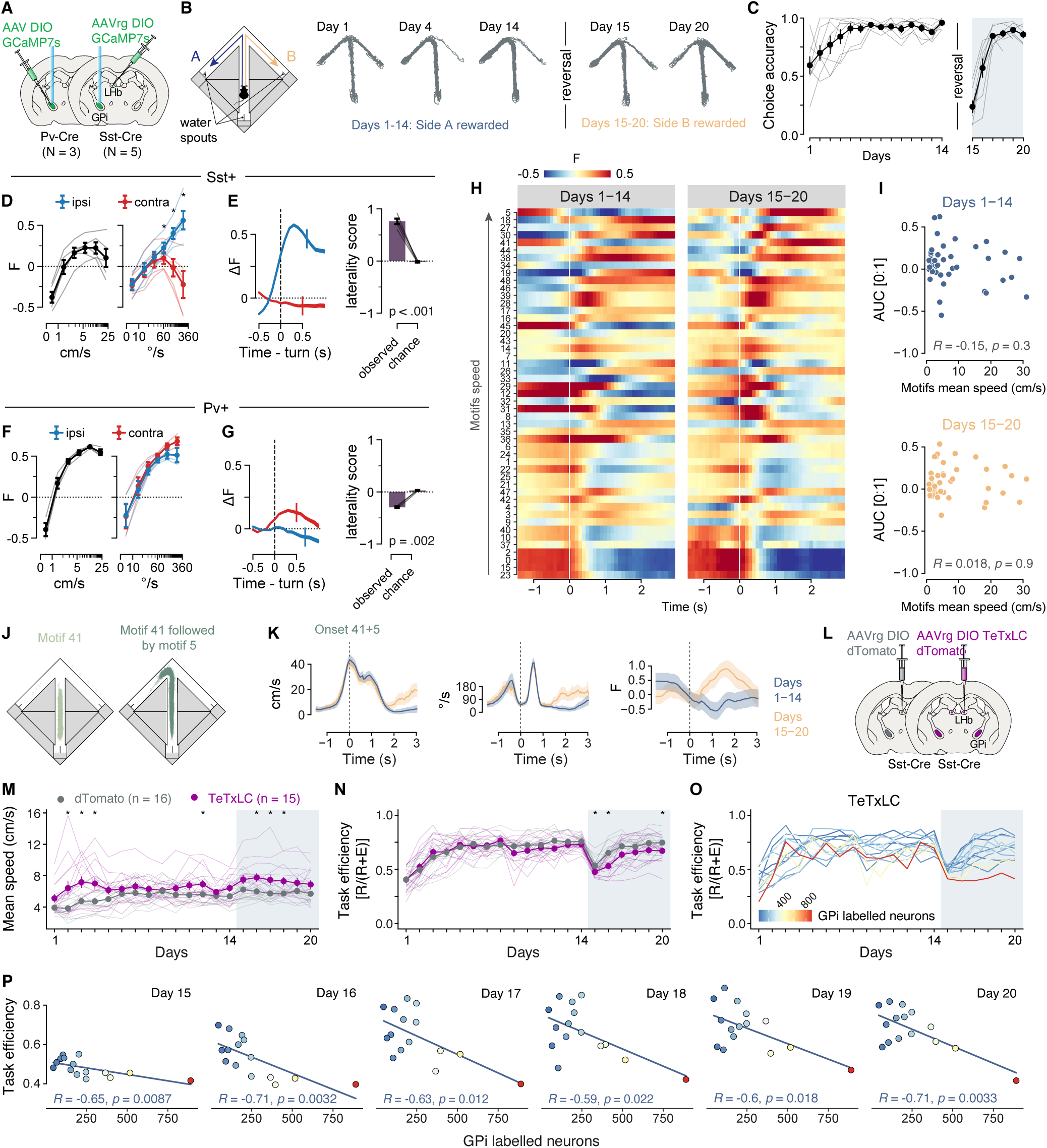
Changes in Sst+ GPi-LHb activity during a maze choice task. **A**, Schematic representation of the experimental strategy: AAVrg-DIO-jGCaMP7s was injected unilaterally in the LHb of Sst-Cre mice (N = 5). AAV-DIO-jGCaMP7s was injected unilaterally in the GPi of Pv-Cre mice (N = 3). 200μm fibers were implanted above the GPi. See also **Supplementary Figure 7A-B** and **Supplementary Table 2**. **B**, Illustration showing the single-reversal maze task. For 14 days the choice to turn contraversive relative to the imaged hemisphere was reinforced with a water drop (13 μl) at the end of the corridor (blue arrow). On day 15 and thereafter, the rewarded choice was reversed (orange arrow). Example traces of one mouse’s paths through the maze are reported for days 1, 4, 14, 15 and 20. See **Methods** for details. **C,** Choice accuracy of the fiber photometry-recorded mice in the single-reversal maze task across days. Gray lines: individual Sst-Cre and Pv-Cre mice. Circles: mean ± SEM. N = 8 mice. **D**, Population activity (F) at different velocities (left) and angular velocities split by turn direction (right) in the open field. Bins spaced evenly along the log-scaled x-axes. Thin lines: individual Sst- Cre mice. Circles: mean ± SEM. *: p < 0.05; paired t-tests, Holm-Sidak correction. N = 5 mice. **E**, Left: ΔF during task-engaged ipsi- and contraversive turns, aligned to turn onset. Baseline for ΔF: mean F of the 0.5s preceding the onset of the turn. Traces: mean ± SEM. Vertical lines: median turn offset times. Ipsi/contra n = 3031/2901 turns (pooled, N = 5 mice). Right: ROC-AUC score-based measure indicating how well single-turn mean ΔF discriminates turn direction. Positive vs negative scores indicate higher activation during ipsi- vs contraversive turns. Chance level established by shuffling the turn direction labels prior to score computation. Gray lines: individual Sst-Cre mice. Bars: mean ± SEM. p < 0.001; paired t-test. N = 5 mice. **F-G**, same as D-E for Pv-Cre mice. **F**, all bins p > 0.2; paired t-tests, Holm-Sidak correction. N = 3 mice. **G**, Left: Ipsi/contra n = 1601/1628 turns (pooled, N = 3 mice). Right: p = 0.002; paired t-test. N = 3 mice. **H**, Mean average of calcium transients of Sst+ GPi-LHb neurons aligned to motifs onset during the pre- and post-reversal phase. The motifs were ordered according to the motif’s speed. **I**, Area under the curve (AUC) of the signal (period between 0 and +1 sec) correlated with the motif’s speed during the pre- and post-reversal phase. Spearman’s correlation coefficients are reported. **J**, Traces of the movements of the mice during motif 41 (in light green, “straight run”) and 41 followed by 5 (in dark green, “contraversive turn”) in the maze arena. **K**, Calcium transients of Sst+ GPi-LHb neurons, angular and linear speed of recorded mice aligned on the end of motif 41 and the beginning of motif 5 during the pre- and post-reversal phase. Mean ± SEM. **L**, AAVrg-DIO-dTomato (N = 16) or AAVrg-DIO-TeTxLC-dTomato (N = 15) were injected bilaterally in the LHb of Sst-Cre mice (see also **Supplementary Figure 7C**). **M**, Mean speed and **N,** task efficiency of AAVrg-DIO-dTomato and AAVrg-DIO-TeTxLC-dTomato injected mice in the single-reversal maze task across 20 days of test. Mean ± SEM. *: p < 0.05, multiple comparisons using the Mann-Whitney U test followed by Holm-Sidak correction. **O**, AAVrg-DIO-TeTxLC-dTomato injected mice’s task efficiency across days color-coded with the number of TeTxLC-expressing cells into the GPi. **P**, AAVrg-DIO-TeTxLC-dTomato injected mice’s task efficiency during post-reversal days regressed on the number of TeTxLC-expressing cells. Pearson’s correlation coefficients are reported. See **Supplementary Table 3** for more information about statistical tests. Abbreviations: F: fluorescence, AUC: area under curve, R: rewards, E: errors, GPi: globus pallidus interna.

We again investigated the GPi activity during rewarded and non-rewarded outcomes to measure negative valence or inverse reward prediction error signaling (**Supplementary Figures 9A**). We found post-reversal Sst+ GPi-LHb activity incompatible with either evaluation signal. If the Sst+ population transmitted inverse reward prediction error, activity should markedly decrease in response to rewards and increase in response to non-reward outcomes following the reversal. Instead, session-average outcome-aligned Sst+ activity remained stable throughout the experiment following rewards and slightly decreased (rather than increased) following non-reward outcomes post-reversal (**Supplementary Figure 9B-C**). Notably, this post-reversal decrease was once again explained by the turning preference: in the maze, mice tended to make contraversive turns after failing to obtain rewards in the contra choice corridor end zone, and contraversive turning was associated with decreased Sst+ GPi-LHb activity as was observed in the serial reversal task (**Supplementary Figures 9B-C**; also see **Figure 4F**, ipsi port). Critically, any putative reward and non-reward modulation of the Sst+ GPi activity remained similar in magnitude during the entire task, arguing against a reward prediction error signaling (**Supplementary Figures 9B-C**). Analysis of the Pv+ GPi activity in reward and non-reward outcomes showed the same pattern as in the serial reversal task (**Supplementary Figures 9D-E**). These findings further support our conclusion that Sst+ GPi-LHb activity does not encode negative valence or prediction error signals.

To test the hypothesis that Sst+ GPi activity encoded the motivational or goal-directed dimension of specific motor actions, we compared the relationship of GPi subpopulation activities with the kinematic profiles across motivational contexts. To do so, we categorized motor motifs in the serial reversal task, and we again found that Sst+ GPi-LHb population activity showed a preference for ipsiversive turns (**Figures 5D-E**). In comparison, Pv+ GPi neurons displayed increased activity during contraversive turns, although the activity amplitude during turning was less pronounced than in the open field (**Figures 5F-G**). Using unsupervised classification, we segmented the behavior into discrete motifs using VAME. When we ranked motifs by their velocity, we surprisingly observed that the activity of Sst+ GPi-LHb neurons showed no relation to the velocity or vigor of action motifs, which was in stark contrast to our observation in the open field (**Figures 5H-I**). These findings further support that the action-specific modulation of Sst+ GPi-LHb activity is defined by the motivational or task context.

We then investigated whether the observed action and sequence-specific modulation of Sst+ GPi-LHb neurons changed following task reversal, focusing on the key choice period captured by the motif sequences in the central arm that described straight running and contraversive turning (i.e., motif 41 “straight run” followed by motif 5 “contraversive turn”, **Figure 5H-J** and **Supplementary Figure 9F** and **10A**). When we aligned calcium signals to the onset of motif 41 followed by motif 5, we found that Sst+ GPi-LHb neurons showed a significantly increased activity after task reversal even if mice performed motor actions at comparable linear and angular speeds during pre- and post-reversal (**Figure 5K**). Notably, the contraversive turn was the rewarded correct choice before the task reversal, whereas it became the incorrect, unrewarded action after the reversal. The observed increase in activity specifically during this transition suggests that Sst+ GPi-LHb neurons contribute to action reversal or behavioral flexibility by signaling when an action that was previously rewarded must be updated to fit the new task contingencies.

To investigate whether Sst+ GPi-LHb neurons play a role in shifting a persistent choice to an alternative option and to establish a potential causal link between neuronal activity and behavior, we conducted cell-type specific silencing experiments in the reversal task. Specifically, we inactivated Sst+ GPi neurons by bilaterally injecting a retrograde AAV with Cre-dependent expression of the tetanus light chain (TeTxLC) into the LHb of Sst-Cre mice (N = 15, **Figure 5L** and **Supplementary Figure 7C**). We compared this experimental group with a control cohort (N = 16) injected with a AAV expressing dTomato. We used the single-reversal maze task to study the effects of Sst+ GPi pathway silencing, and similarly to the motor-related changes in the open field (**Figure 4O**) we found an increased motif speed and reduced immobility time (**Figure 5M** and **Supplementary Figure 11B**). The cell-type specific silencing also increased the number of trials performed by the Sst+ GPi TeTxLC mice, in particular during the first training days and immediately after the reversal. During the pre-reversal phase, mice with TeTxLC silenced Sst+ GPi-LHb neurons reached accurate and asymptotic performance (**Figure 5N** and **Supplementary Figures 11C-E**). However, in the post-reversal phase, we found that silencing Sst+ GPi-LHb neurons resulted in an increased number of error trials (**Supplementary Figures 11A, E, D**). As a result, mice with silenced Sst+ GPi-LHb neurons showed a decline in task efficiency during the first days after reversal indicating an impairment in the ability to adjust their choice strategy and suppress the well-trained pre-reversal action response (**Figure 5N**). Importantly, after reversal we observed that TeTxLC mice showed increased variability in performance, and their impaired behavioral efficiency was strongly correlated with the number of Sst+ GPi-LHb neurons expressing TeTxLC (**Figure 5O-P**). These findings demonstrate that Sst+ GPi-LHb neurons play a central role in adjusting goal-directed behavior and driving choice flexibility, through suppressing well-trained actions and promoting the alternative choice after task reversal.

### Flexibility-driven modulation of Sst+ GPi-LHb activity in a single-reversal maze task

As the most prominent activation of Sst+ GPi-LHb neurons occurred in the central arm and entering the choice junction, we further analyzed the neuronal activity during this behavioral trajectory (**Figure 6A**). Mice naturally showed a decline in motivation and task-engagement in the later phases of the session (**Figure 6A** and **Supplementary Figures 9G-I**), and we therefore divided trials into four quartiles (Q1-4) to assess how neuronal activity changed with the motivational state. We found that Sst+ GPi-LHb activity became suppressed as mice performed choice runs with contraversive turning early in the session (i.e. Q1, Q2), while activity increased during the same trajectory in the last trials as mice decreased their velocity (i.e. Q4; **Figure 6B**). In contrast, in sessions after reversal of the reward location (i.e. post-reversal sessions), the Sst+ GPi-LHb activity during the same non-rewarded trajectory increased even in early trials (**Figure 6B**). Similar to our findings in the serial reversal task, Sst+ GPi-LHb neurons were more active during choice runs with ipsiversive turning, and this ipsiversive turning related activity consistently increased as mice approached the turn across all session quartiles (**Figure 6B**).

**Figure 6.**
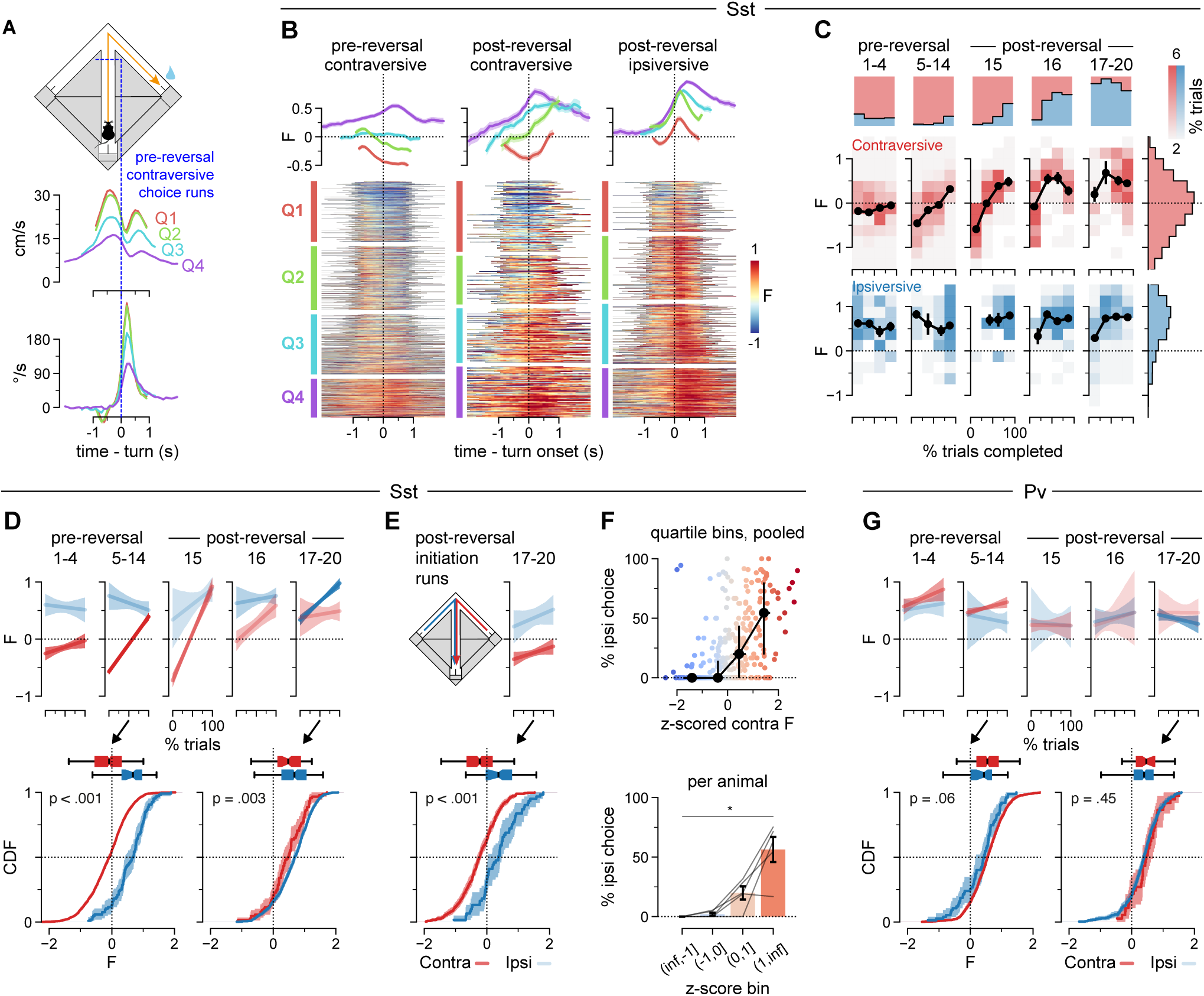
Dynamic modulation of Sst+ GPi-LHb neuronal activity during strategy shifts. **A**, Average velocities (middle) and contraversive angular velocities (bottom) of Sst-Cre mice during pre-reversal contraversive choice runs in each session-trial quartile bin (Q1-4, see also panel **B**), aligned to turn entry (top). Traces are clipped at the median initiation zone exit (run start) and reward zone entry (run end) time points. Data from sessions 1-4 excluded. Traces: mean ± SEM. Q1-4 n = 424/446/418/271 turns (pooled, N = 5 mice). **B**, Raster: Sst+ GPi-LHb population activity (F) for individual pre- and post-reversal choice runs. Trials were grouped by task stage and turn direction, sorted by their relative position in their session (i.e., the percentage of session trials completed; trial number/session total), and finally binned into quartiles (Q1-4). The bins are unevenly sized due to the exclusion of trials in which mice returned to the center corridor before reaching a reward zone. Data from sessions 1-4 and pre-reversal ipsiversive trials omitted, the latter due to the low number of trials available. Traces: mean ± SEM for each raster bin; clipped as in panel A. n = 1559/397/826 trials (pooled, N = 5 mice). **C**, Top row: stacked histograms indicating the fraction of contra- vs ipsiversive choices, by session quarter, over the course of the experiment. Quarters are based on trial quartiles, not time. Trials pooled over Sst-Cre mice. Middle and bottom rows: density plots visualizing Sst+ activity (F) during contra- (middle, in red) and ipsiversive (bottom, in blue) choice turns, by session quarter, over the course of the experiment (same as above). One data point, mean F, included per turn. Circles: mean ± SEM of turns by session quarter; mean omitted for bins with fewer than 5 data points total. Right-hand histograms: distribution of mean turn F pooled over all sessions. Contra/ipsi n = 2416/1013 trials (pooled, N = 5 mice; cf. panel D). **D**, Top row: Linear regression analysis predicting single-trial Sst+ GPi-LHb choice turn activity based on session progress (i.e., the percentage of session trials completed; trial number/session total) and turn direction. Lines: regression fits and 95% confidence intervals (bootstrapped); increased transparency indicates smaller data sets. Contra n = 460/1559/216/93/88 and ipsi n = 82/105/45/107/674 (pooled, N = 5 mice). Bottom row: distributions of Sst+ GPi-LHb activity (F) during choice turns, plotted by turn direction, for session bins “5-14” (left) and “17-20” (right). Boxplots: center lines represent the distribution median; edges, the upper and lower quartiles; whiskers, the 2.5% and 97.5% points of the distribution. Lines: cumulative distribution functions (CDFs) and 95% confidence intervals (bootstrapped). The CDF indicates the fraction of turns during which F was at or below a given value. p-values: Kolmogorov-Smirnov test, Holm-Sidak correction. “5-14” bin contra/ipsi n = 1559/105 and “7-20” bin contra/ipsi n = 88/674 (pooled, N = 5 mice). **E**, Plots as described in D, but for post-reversal “return” runs from the choice zones towards the initiation zone (days 17-20). Bottom panel p-value computed and corrected with those in D. Contra/ipsi n = 369/61 (pooled, N = 5 mice). **F**, Top: probability of ipsiversive choices plotted against the average F during contraversive turns for all session-quartile bins. Small circles: individual session-quartile averages, contra turn F z-scored per mouse. Black circles: median and IQR of binned session-quartiles; bin edges as indicated in bottom panel. n = 332 session-quartiles (pooled, N = 5 mice). Bottom: same as above, plotted per mouse. Thin lines: median of binned session-quartiles for individual mice; bin edges are indicated on the x axis. Bars: mean ± SEM. *: p < 0.001, repeated-measures ANOVA. N = 5 mice. **G**, Same as D for Pv-Cre mice. Top row: contra n = 211/841/79/34/37 and ipsi n = 63/75/40/97/345 (pooled, N = 3 mice). Bottom row: “5-14” bin conta/ipsi n = 841/75 and “17-20” bin contra/ipsi n = 37/345 (pooled, N = 3 mice). See **Supplementary Table 3** for more information about statistical tests.

We then explored how the suppressed Sst+ GPi activity during the choice phase was related to task performance over sessions. To examine this, we plotted neuronal activity during the turn against the fraction of completed trials in each session (i.e., trial number/session total). Notably, as mice learned the task and developed a preference for correct contraversive turning, the suppression of Sst+ GPi activity in early trials became more pronounced (**Figures 6C-D**). After task reversal, when mice began to redirect their turning toward the ipsiversive choice corridor (i.e. the new rewarded side), the contraversive turning instead showed positive transients, resulting in similar Sst+ GPi activity during both ipsi- and contraversive turning (**Figures 6C-D**). Remarkably, Sst+ GPi activity during turning to the start zone (i.e., during trial initiation runs) showed no such modulation, as activity during contraversive “return” turns remained moderately suppressed and distinguishable from ipsiversive turn-related signals (**Figures 6E**). This indicates that the loss of the contraversive choice-related suppression is choice-specific and likely attributable to reversal learning.

To further investigate the relationship between Sst+ GPi-LHb neuron activity and choice behavior, we binned contraversive turning trials by session and quartile and averaged the neuronal activity. When plotting these averages against the percentage of ipsiversive choices within the same bin, we observed that higher activity during contraversive trials correlated with greater ipsiversive choice probability (**Figure 6F**). This suggests that elevated activity in Sst+ GPi- LHb neurons during contraversive trials, and by extension a reduction in directional selectivity of the Sst+ neurons, reflect a state more permissive of ipsiversive turning. In other words, whereas low Sst+ activity preceding a turn reliably signals a contraversive turn, high activity may be followed by turns in either direction, likely depending on the level of suppression in the opposite hemisphere. Unlike Sst+ GPi-LHb neurons, the Pv+ GPi population maintained consistently high activity levels during both contraversive and ipsiversive runs, showing little sensitivity to turn direction, task-engagement, or reversal condition (**Figures 6G** and **Supplementary Figure 12A-C**). In summary, we found that the activity of Sst+ GPi neurons during the choice phase reflects an updating of decision signals related to the reward location, which supports their role in adjusting goal-directed behavior and choice flexibility.

## Discussion

The basal ganglia play a central role in selecting and evaluating actions, functions that are mediated via parallel but distinct GPi output pathways^12,15,16^. In this framework, actions are selected by GPi neurons that project to motor thalamus and brainstem, and action outcomes are evaluated by GPi neurons that project to the LHb^12,16^. Here, we have expanded the anatomical and functional characterizations of the GPi subtype that is characterized by projections to the LHb (GPi-LHb subtype) to define their role in goal-directed behavior. We report that the main GPi-LHb pathway – genetically defined by Vgat/Vglut2/Sst co-expression (Sst+ GPi)^13,15^ – possesses extensive collateral projections that target the GPe, the striatal striosomes, as well as dopamine neurons in the SNc. Our data further support that Sst+ GPi neurons co-release GABA and glutamate in all these target regions, similar to their co-release profile in the LHb^13,14,27^. In light of this unique neuroanatomical organization, which positions Sst+ GPi neurons to broadcast a wide modulatory feedback signal in the basal ganglia, we decided to investigate their role in shaping learning and execution of choice actions.

The GPi projections to LHb have been proposed to contribute to inverse reward prediction errors and reinforcement learning through a LHb activity-mediated suppression of dopamine release in the basal ganglia action selection circuit ^12,16,28–30^. However, recent studies indicate that the negative valence and reinforcement learning signals found in the LHb originate – to a functionally significant degree – from excitatory inputs from the LHA^13,31^ and other regions^18,32,33^, rather than from the GPi. Importantly, our findings showing that Sst+ GPi neurons do not encode inverse reward prediction errors nor negative outcome value agrees with results of precise optogenetic activation of Sst+ GPi neurons and their LHb projections at the time of the trial outcome that fails to change decision-making^13^.

As highlighted in prior studies, the LHb plays a pivotal role in processing aversive signals^34,35^, primarily by increasing activity in response to negative outcomes, which suppresses midbrain dopamine release^28,36^. This function is critical for guiding adaptive behaviors, as LHb inactivation impairs outcome-based decision-making^37^, including effort-based choices^38^ and adaptive avoidance learning^39^. While our data suggests that Sst+ GPi neurons do not directly encode negative valence, their projections to the LHb may instead modulate the activity of LHb circuits and their specialized outputs involved in adaptive behaviors. This hypothesis aligns with the growing evidence of functional heterogeneity within the habenula and its complex connectivity^40^. We propose that distinct subpopulations of LHb neurons and their discretely organized inputs may mediate these dual or more complex roles of the LHb, with Sst+ GPi inputs shaping the flexibility of behavioral outputs without directly engaging in aversive signal processing.

A key finding of our study is the context-dependent activity of the Sst+ GPi pathway. We show that Sst+ GPi activity reflected the turn direction in mice that performed a spatial reversal task for water rewards, more so than when they spontaneously explored a reward-free environment. Specifically, the Sst+ GPi population activity was strongly lateralized and correlated with ipsiversive turning choices in the reversal tasks, and this activity profile was absent during exploration of an open field without reinforcement. Similar examples of reward or task context-dependent modulations of movement-related signals have previously been found in GPi neurons^41–43^. In comparison, the Pv+ GPi population displayed the opposite pattern, as Pv+ GPi neurons were strongly activated during contraversive turning in the open field but not in the choice tasks. Our findings on the task-specific activity of Sst+ GPi neurons, showing their activation during choice turning direction but not outcome-related prediction errors, supports their role in progressively shaping action selection instead of immediate action evaluation. This distinction is supported by our experiments showing that inactivation of Sst+ GPi neurons (i) increases overall locomotor activity without impairing motor execution per se, yet (ii) degrades optimal performance in the single reversal task. Notably, studies have shown that LHb lesions or inhibition produce similar effects on locomotion and reversal performance^38,39,44^.

The activity profile of Sst+ GPI neurons and the behavioral effects following their genetic silencing together support an important role in the development of a choice-turning activity that evolves with training during task learning. Specifically, we found that the Sst+ GPi activity during contraversive choice turning gradually increased after reversal of the reward location, whereas the activity recorded during ipsiversive choice turning remained stationary. As a result, the lateralized turning-selective activity gradually decreased as mice learned to shift their turning preference from contra- to ipsiversive actions. Interestingly, the striatum also shows population-level activation during contraversive choices^45^. We propose that this gradual rebalancing of contraversive and ipsiversive choice-related activity during adaptation to a new task rule reflects the key role of the Sst+ GPi population in suppressing previously learned and preferred actions. Here, we have shown how the organization of the Sst+ GPi pathway expands the complexity of connectivity diagrams that depict the basal ganglia circuit as a set of one-directional parallel pathways. Our results show that the genetically defined Sst+ GPi population forms an unexpected feedback projection in the basal ganglia, and we propose that this organization can underline a neuromodulatory signal that suppresses the commitment to repeat the same action, thereby promoting the exploration of alternative actions in the environment.

## Acknowledgements

We thank the Animal Behavior Core Facility (ABCF) at Karolinska Institutet for access to infrastructure and Alexander Wolthon for mouse colony management. Funding for this study was provided to K.M. by the Swedish Research Council (Vetenskapsrådet MH), the Swedish Brain Foundation (Hjärnfonden), Karolinska Institutet (KID doctoral funding for V.S. and M.W.), and the Knut och Alice Wallenberg Foundation. Additional funding was provided by the Swedish Society for Medical Research (Svenska Sällskapet för Medicinsk Forskning) and the Wenner Gren Foundation to S.Ä-R.; the Swiss National Science Foundation (project number: P500PB_214355) and the Amicitia Foundation to A.C.

## Author Contributions

SÄ-R, VS and IL designed, interpreted and performed the anatomical characterization. SÄ-R and VS analyzed and visualized the anatomical data with the help of IS and AC. MW and IL designed the behavioral experiments. MW, IL, AC and VS performed the behavioral experiments with the help of IS. MW and AC analyzed and visualized all behavioral data. JF performed all electrophysiology experiments and analysis. MW, SÄ-R, VS, AC and K.M. wrote the manuscript. IL and KM supervised and conceptualized the work. KM was responsible for project funding and management. All of the authors discussed and commented on the manuscript.

## Competing Interests statement

The authors declare no competing interests.

## Data availability

All data presented in this paper is available upon request to the corresponding authors.

## Code availability

Custom code is available upon request.

## Notes

### Competing Interest Statement

The authors have declared no competing interest.

### Summary of Updates

Revised manuscript with new data and updated figures.

