## supplementary Figures for "Sst+ GPi output neurons provide direct feedback to key nodes of the basal ganglia and drive behavioral flexibility"

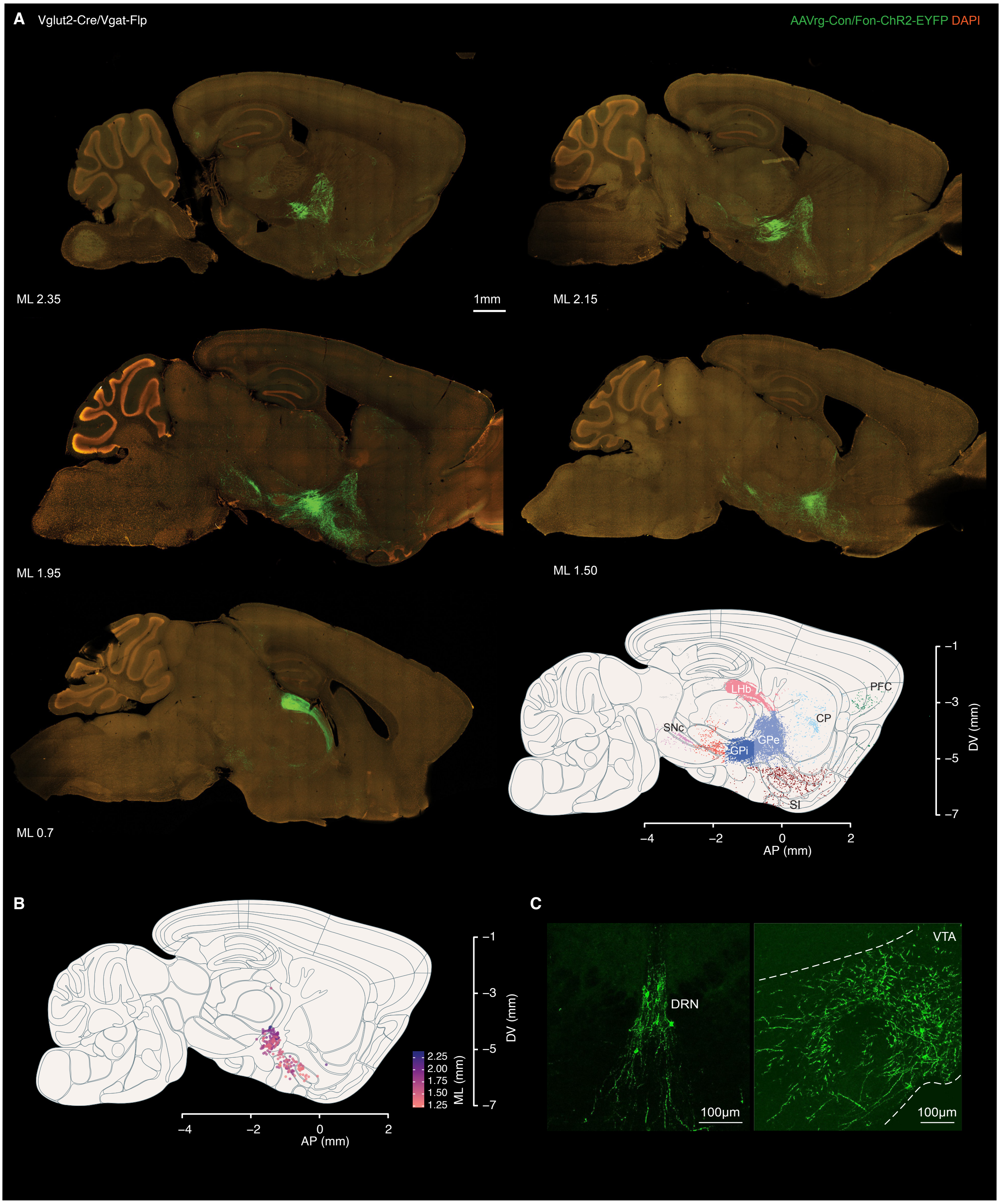

Supplementary Figure 1

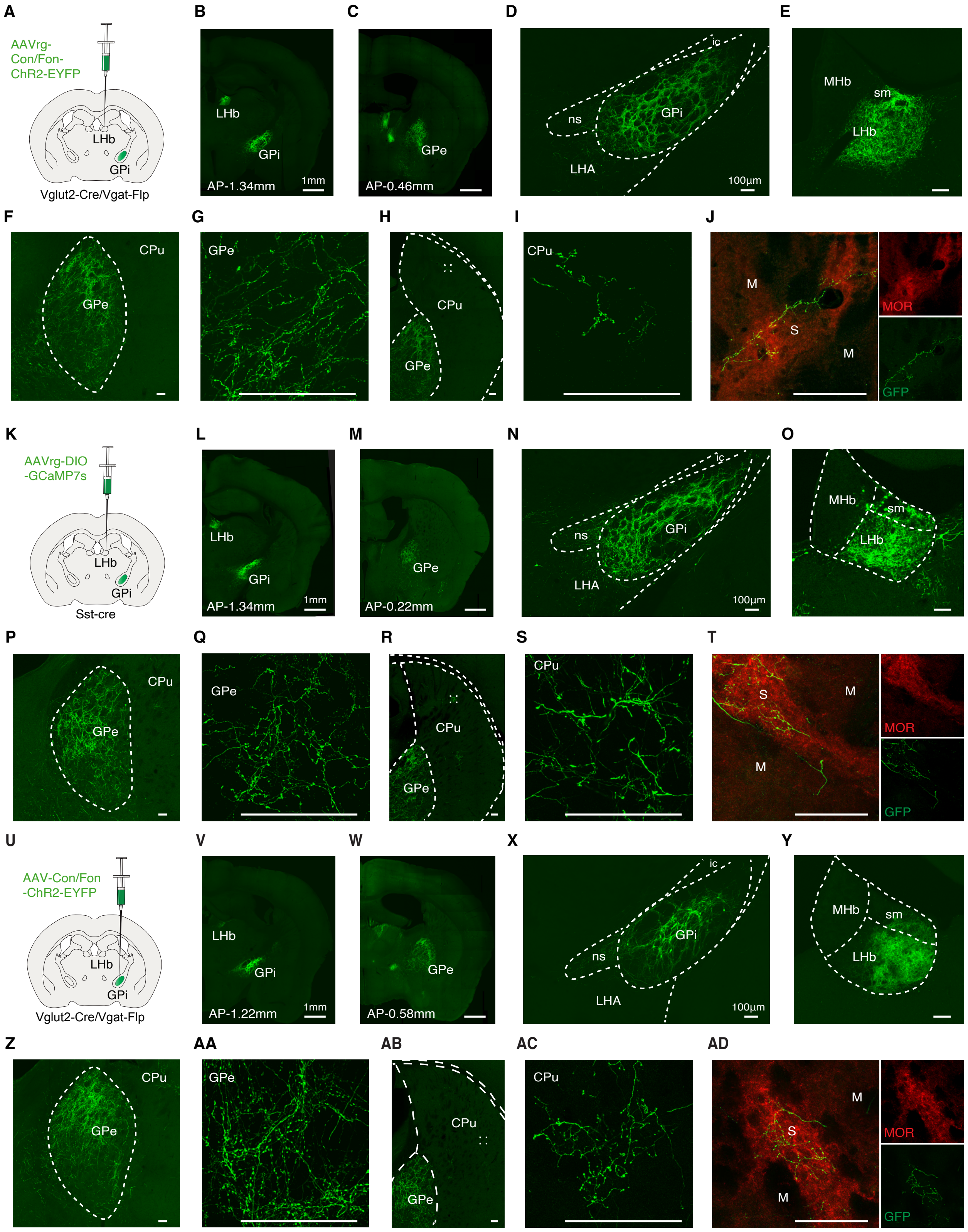

Supplementary Figure 2

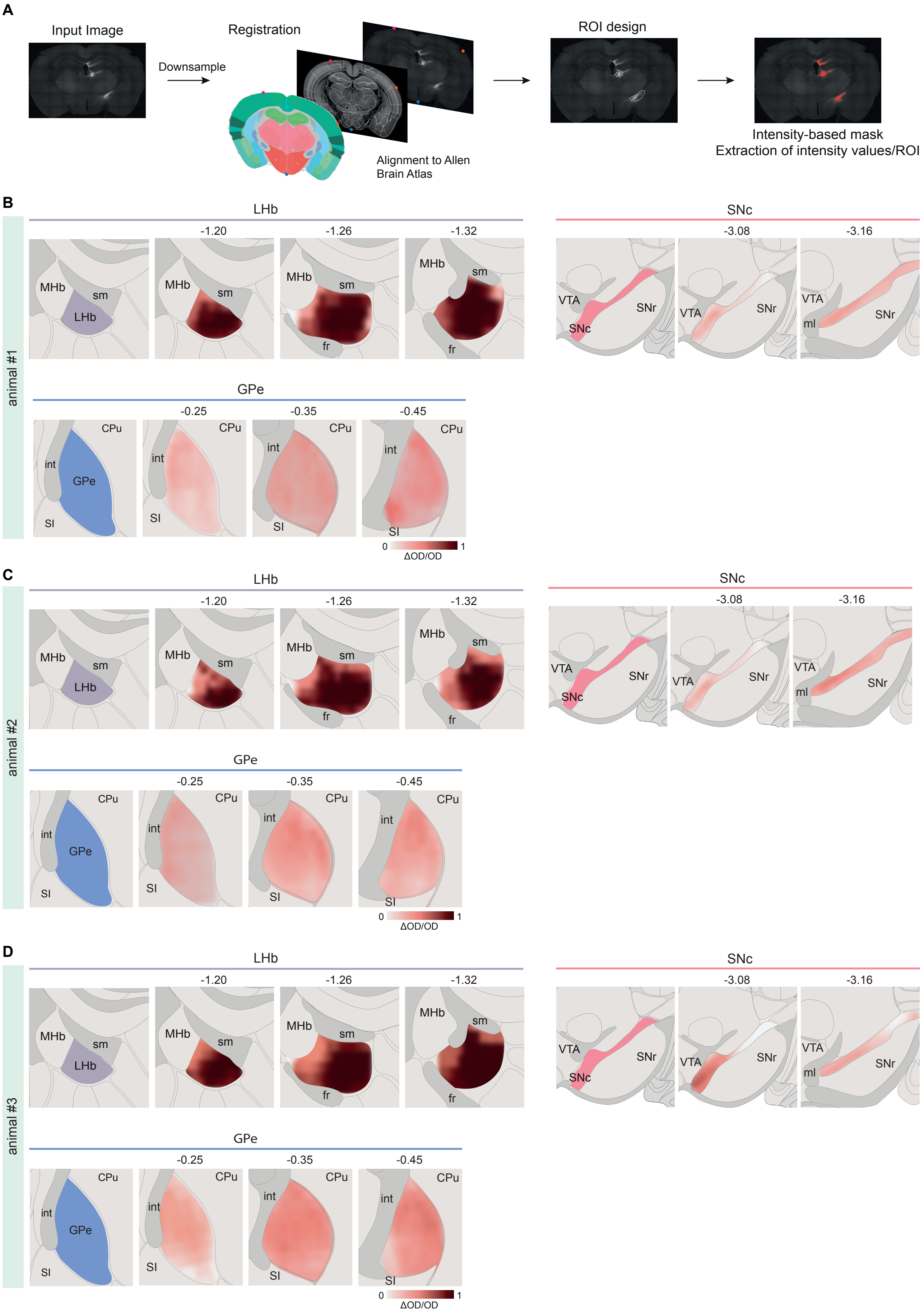

Supplementary Figure 3

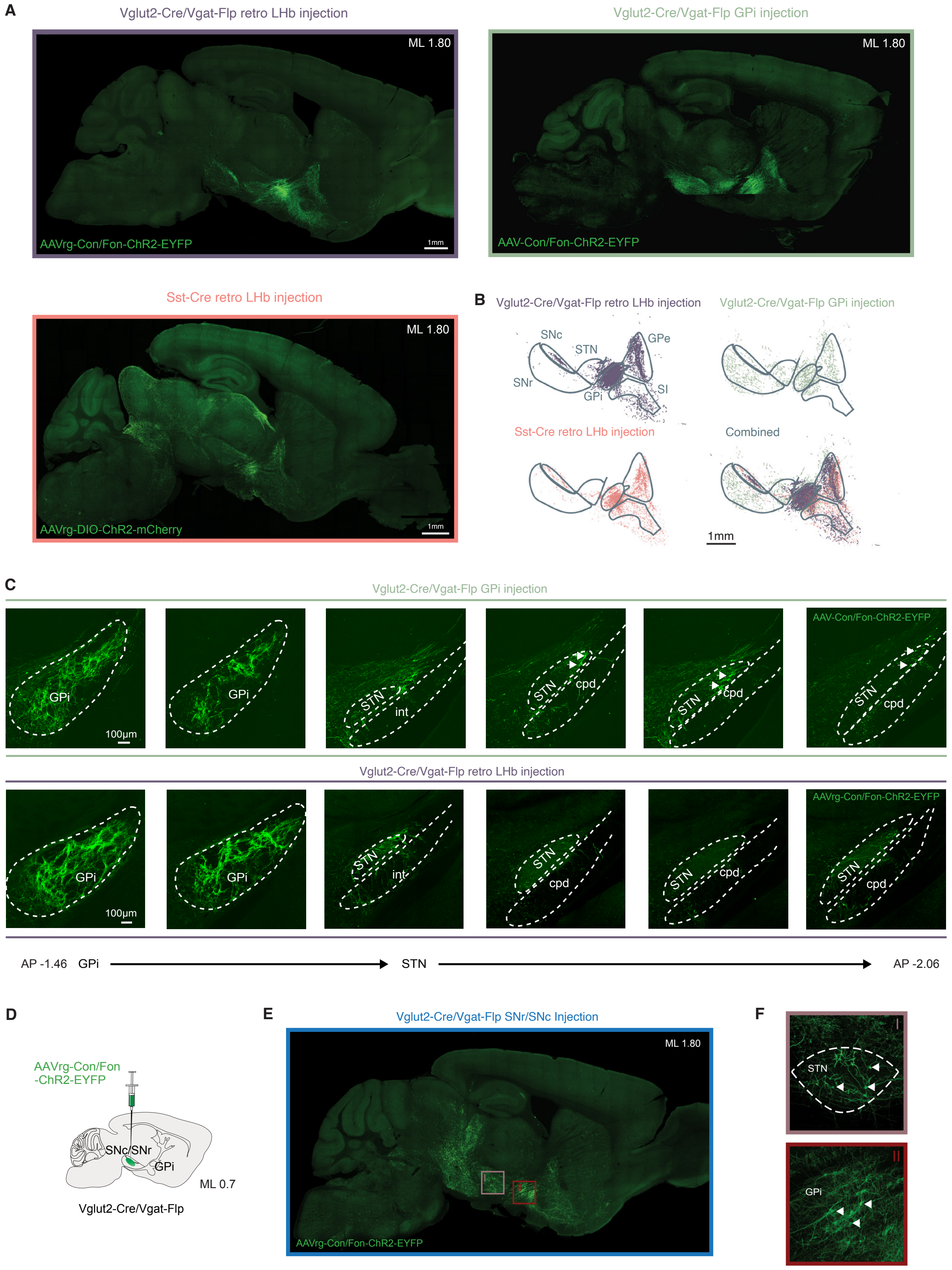

Supplementary Figure 4

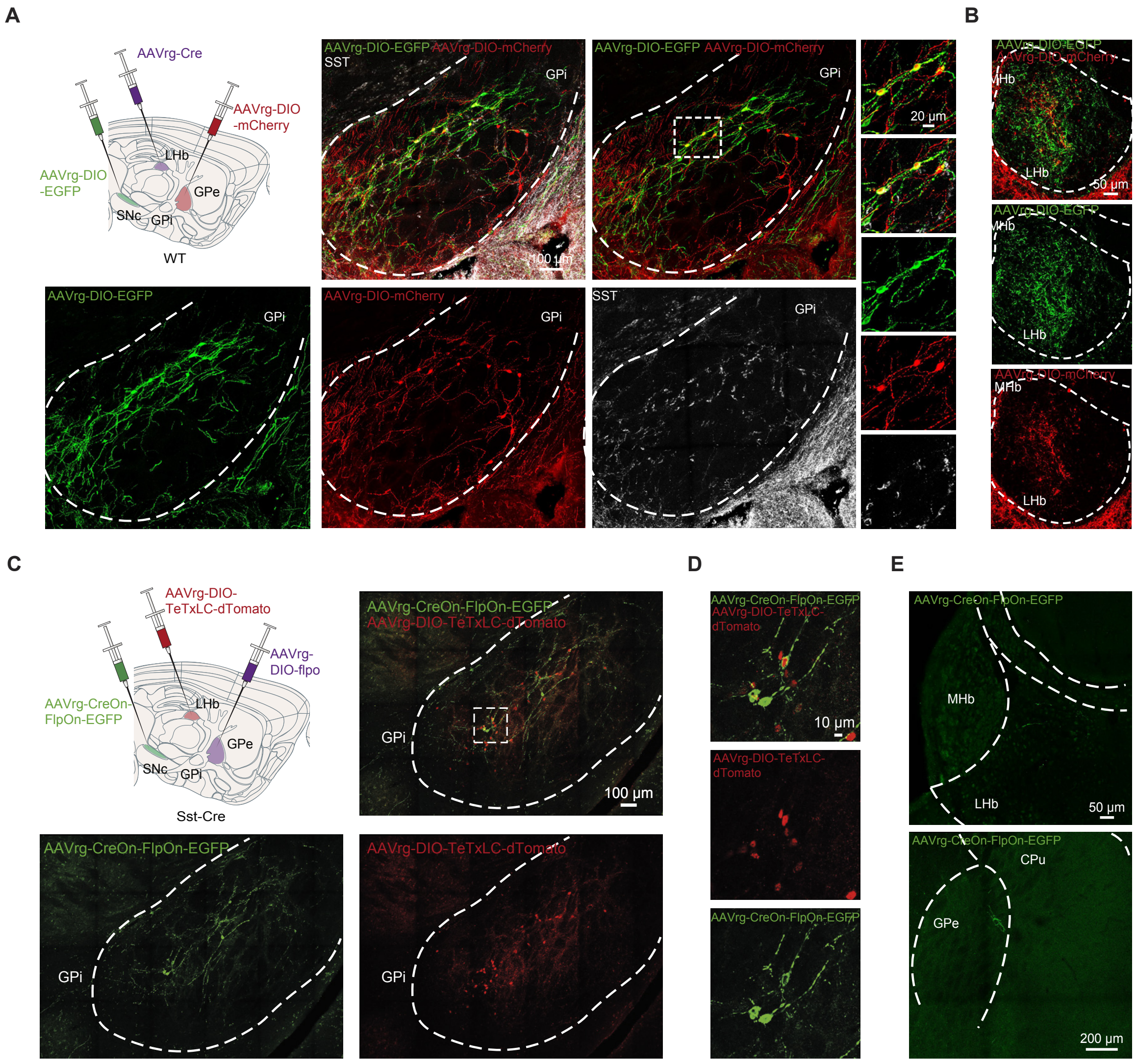

Supplementary Figure 5

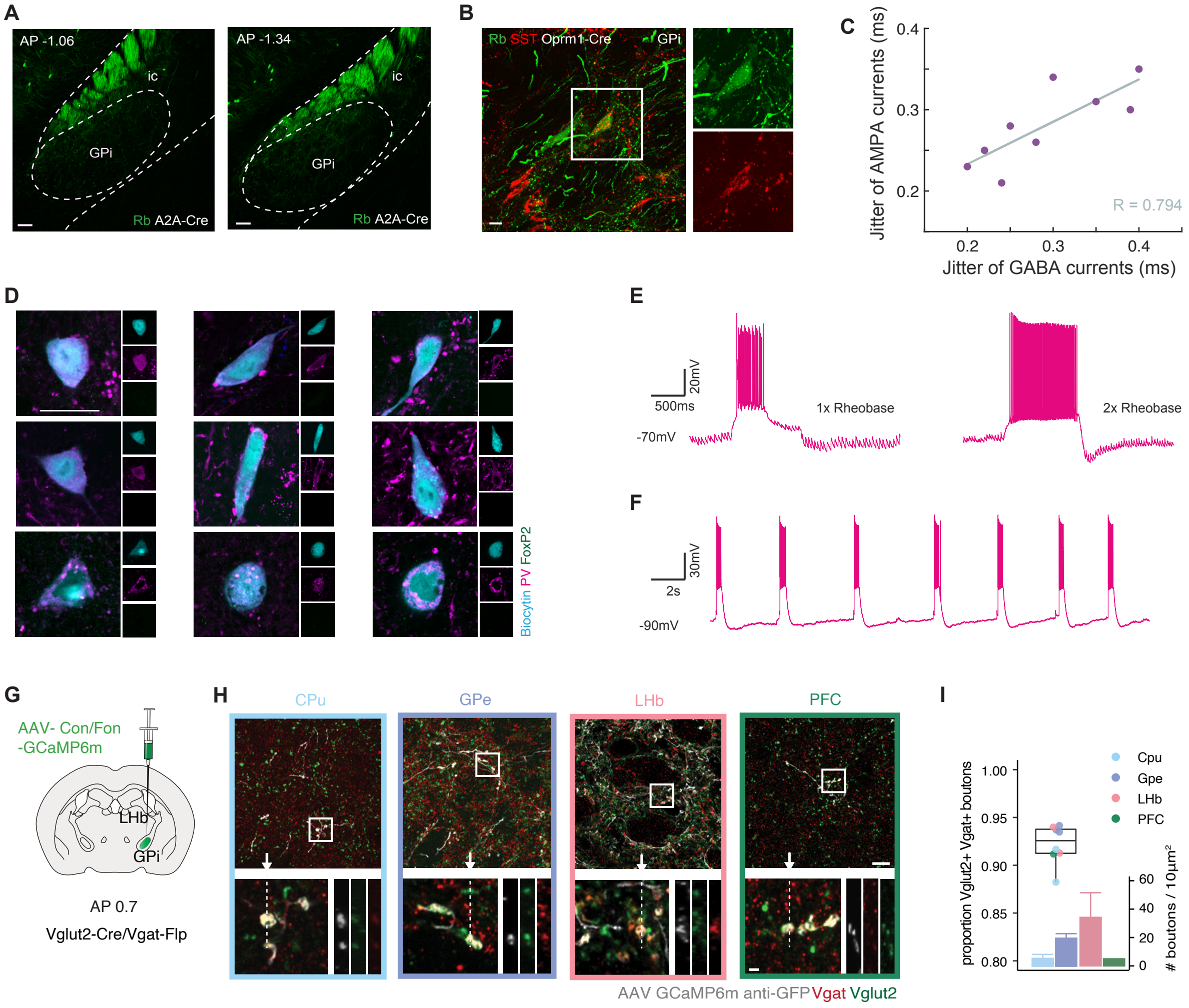

Supplementary Figure 6

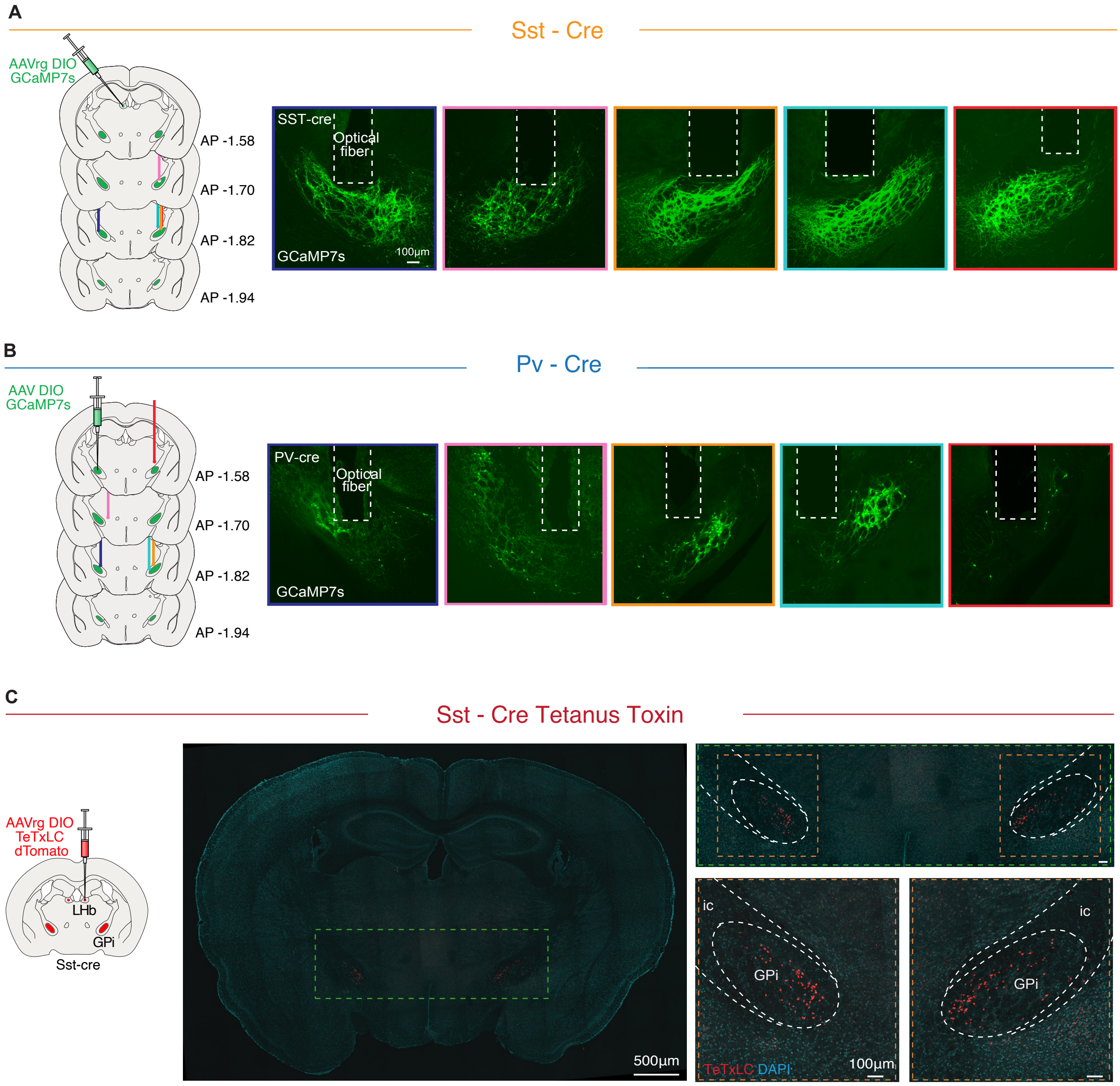

Supplementary Figure 7

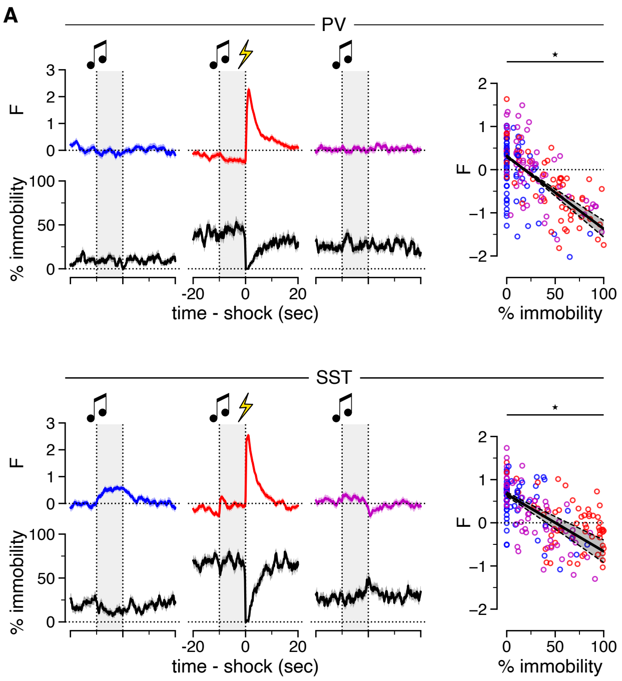

Supplementary Figure 8

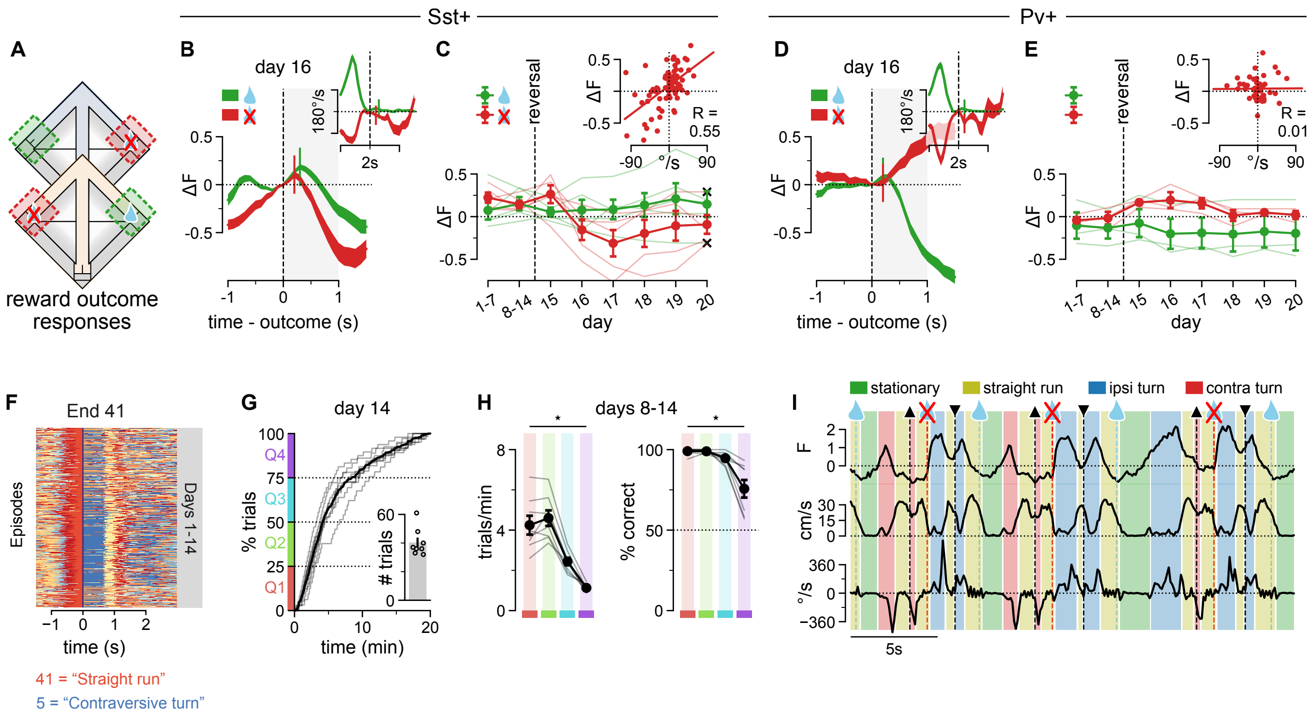

Supplementary Figure 9

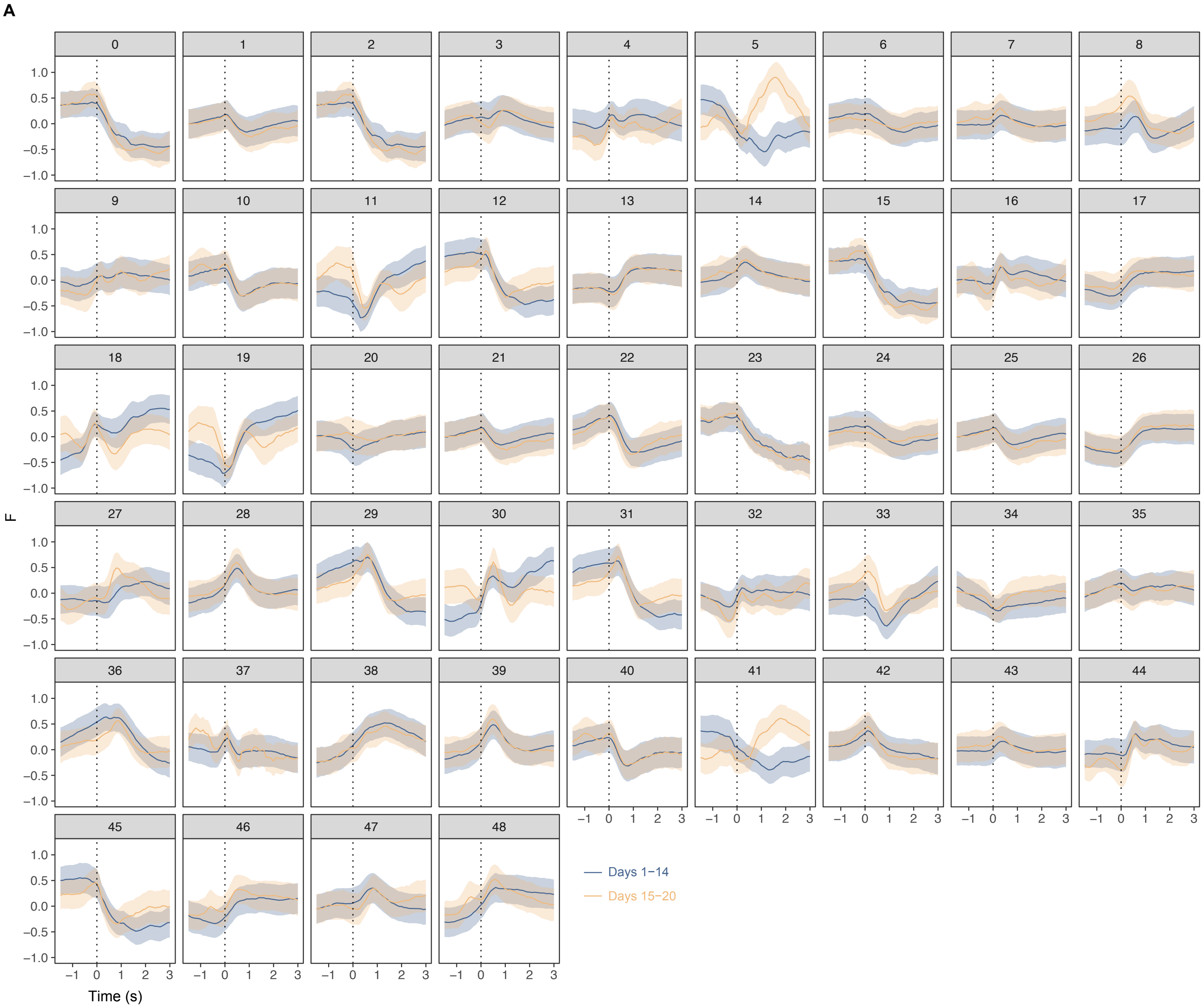

Supplementary Figure 10

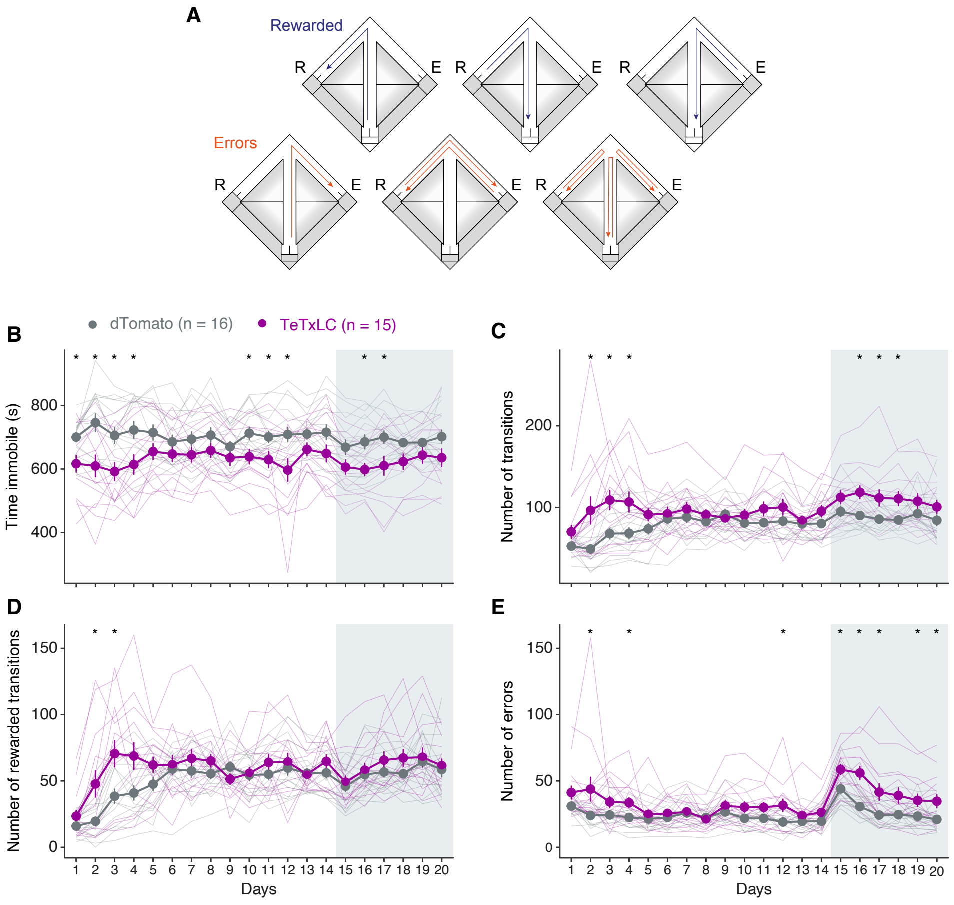

Supplementary Figure 11

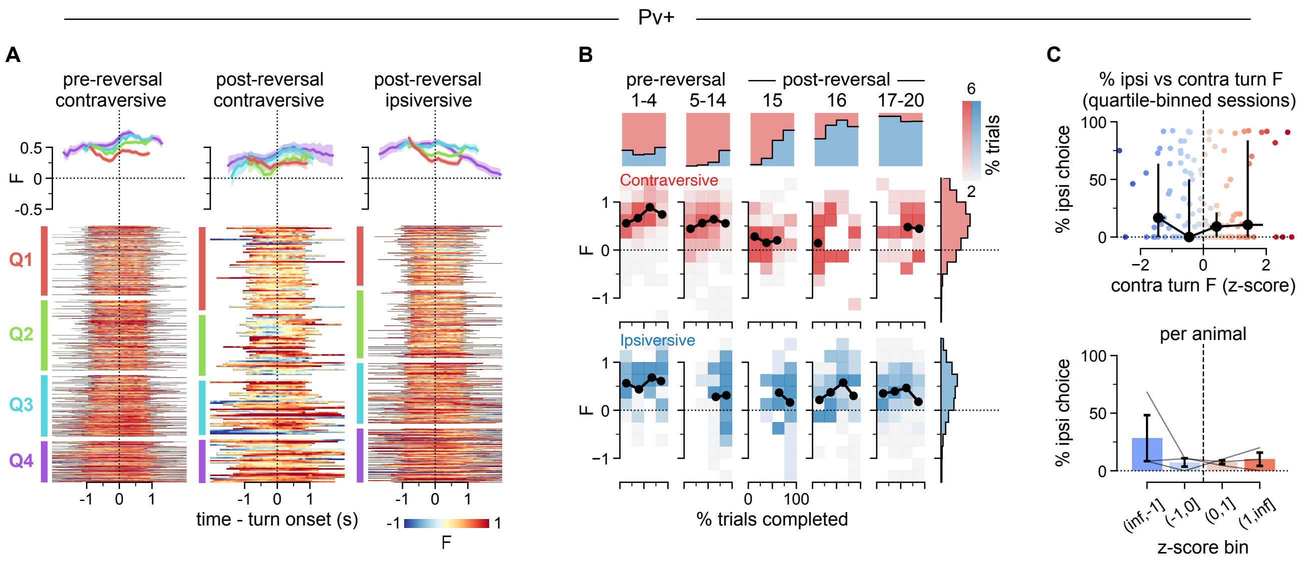

Supplementary Figure 12

**Supplementary Figure 1 | Vglut2+/Vgat+ GPi LHb-projecting neurons can be identified in the GPi, SI, VTA and DRN.**

**A**, Representative sagittal sections from one hemisphere showing AAVrg-ChR2-EGFP-expressing Vglut2+/Vgat+$^{GPi\text{-}LHb}$ axons throughout the hemisphere. Bottom right: Vglut2+/Vgat+GPi-LHb axons throughout the brain, colored based on their target brain region (as in ARA). Brain sections (n = 15) superimposed onto the ARA plate #100884129 (~1.95 ML).

**B**, Graphical representation of LHb-projecting Vglut2+/Vgat+ EYFP labelled cell bodies, identified in the GPi and the SI. Each dot represents one cell body identified within one hemisphere. All neurons are superimposed onto the ARA plate #100884129 (~1.95 ML), each neuron is color-coded by its ML stereotaxic coordinate (blue: lateral; light pink: medial). n = 21 sections.

**C**, Representative images showing LHb-projecting Vglut2+/Vgat+ EYFP expressing cell bodies in the VTA and DRN.

Abbreviations: AP: anteroposterior, DV: dorsoventral, ML: mediolateral, GPI: Globus pallidus interna, SI: Substantia innominata, LHb: Lateral habenula, SNc: substantia nigra pars compacta, GPe: globus pallidus externa, CP: Caudate putamen, PFC: prefrontal cortex, DRN: dorsal raphe nucleus, VTA: ventral tegmental area.

**Supplementary Figure 2 | LHb-projecting GPi neurons targeted with three different viral strategies show projection targets.**

**A**, Experimental strategy: AAVrg-Con/Fon-ChR2-EYFP was unilaterally injected in the LHb of Vglut2-Cre/Vgat-Flp mice.

**B–J**, Representative coronal sections depicting the main projection targets of retrogradely targeted Vglut2+/Vgat+ GPi neurons after injection of AAVrg-Con/Fon-ChR2-EYFP in LHb in Vglut2-cre/Vgat-Flp mice. N = 4 mice.

**K**, Schematic illustration of the experimental strategy: AAVrg-DIO-jGCaMP7s was unilaterally injected into the LHb of Sst-Cre mice.

**L–T**, Representative coronal sections depicting the main projection targets of retrogradely targeted Sst+ GPi neurons after injection of AAVrg-DIO-jGCaMP7s in LHb in Sst-cre mice. N = 9 mice.

**U**, Schematic illustration of the experimental strategy: AAV-Con/Fon-ChR2-EYFP was unilaterally injected directly into the GPi of Vglut2-Cre/Vgat-Flp mice.

**V–AD**, Representative coronal sections depicting the main projection targets of directly targeted Vglut2+/Vgat+ GPi neurons after injection of AAV-Con/Fon-ChR2-EYFP in the GPi in Vglut2-cre/Vgat-Flp mice. N = 3 mice.

**B, D, E, L, N, O, V, X, Y**, Coronal sections of the injected hemisphere displaying cell bodies in the GPi and axons in the LHb for the three viral strategies, respectively. Scale bars: 1mm (**B, L, V**), close-ups: 100μm (**D, E, N, O, X, Y**).

**C, F, G, M, P, Q, W, Z, AA**, Axonal arborizations in the GPe of GPi LHb-projecting neurons were confirmed for the three viral strategies implemented. Scale bars: 1mm (**C, M, W**), close-ups: 100μm (**F, G, P, Q, Z, AA**).

**H, I, R, S, AB, AC**, GPi LHb-projecting neurons' afferent projections show discrete patches of arborization within the CPu. Close-up images of axonal arborization are indicated with a white square in the overview image. Scale bars: 100μm.

**J, T, AD**, The three viral strategies all show collateral projections of the GPi LHb-projecting neurons (green) in the striosome compartment of the CPu identified with MOR immunostaining (red). Scale bars: 100μm.

Abbreviations; LHA: Lateral Hypothalamus, ns: nigrostriatal bundle, MHb: Medial Habenula, sm: stria medullaris, S: striosome compartment, M: matrix compartment, GPi: globus pallidus interna, LHb: Lateral habenula, GPe: globus pallidus externa, CPu: caudate putamen, MOR: mu opioid receptor.

**Supplementary Figure 3 | Quantification of the Sst+ GPi-LHb axonal projections in target brain regions across individual mice**

**A**, Schematic representation of the pipeline used to quantify the intensity of the projections of Sst+ GPi-LHb axons in the target areas.

**B-D**, Heatmap showing the fluorescence intensity of the projections of Sst+ GPi-LHb axons in three anteroposterior levels of LHb, GPe and two anteroposterior levels of SNc, in each animal used in this strategy.

Abbreviations; ROI: region of interest, LHb: Lateral habenula, MHb: Medial habenula, sm: stria medullaris, fr: fasciculus retroflexus, SNc: substantia nigra pars compacta, SNr: substantia nigra pars reticulata, VTA: ventral tegmental area, ml: medial lemniscus, GPe: globus pallidus externa, CPu: caudate putamen, int: internal capsule, SI: substantia innominata, OD: optical density.

**Supplementary Figure 4 | Direct comparison of three viral strategies targeting LHb-projecting GPi neurons.**

**A**, Representative sagittal section from three viral strategies targeting the Vglut2+/Vgat+ and SST+ GPi LHb-projecting neurons. AAVrg-Con/Fon-ChR2-EYFP unilaterally injected in the LHb of Vglut2-Cre/Vgat-Flp mice (purple), AAV-Con/Fon-ChR2-EYFP unilaterally injected into the GPi of Vglut2-Cre/Vgat-Flp mice (green) and AAVrg-DIO-jGCaMP7s unilaterally injected into the LHb of Sst-Cre mice (pink), respectively. Scale bar: 1mm.

**B**, The Vglut2+/Vgat+ or Sst+ GPi-LHb neurons afferent projections from the sagittal brain sections displayed in **A,** displayed separately and overlaid for a direct comparison of projection targets. The three viral strategies for targeting Vglut2+/Vgat+ or Sst+ GPi-LHb neurons give rise to efferent projections with differential targeting of the SNr, STN and SI. Viral strategy color coded as in **A**.

**C**, A direct injection of AAV8-Con/Fon-ChR2-EYFP into the GPi of Vglut2-Cre/Vgat-Flp mice (green) results in Vglut2+/Vgat+ ChR2-expressing cell bodies within the STN (indicated by white arrows), alas suspected to result in axons in the SNr (as seen in **A**, **B**: green). AAVrg-Con/Fon-ChR2-EYFP injected in the LHb of Vglut2-Cre/Vgat-Flp mice (purple) display no cell bodies in the STN. Scale bar: 100µm.

**D**, Illustration describing the experimental strategy: AAVrg-Con/Fon-ChR2-EYFP was unilaterally injected into the SNr and SNc of Vglut2+/Vgat+ mice.

**E**, Representative sagittal section from a Vglut2+/Vgat+ mouse with a unilateral injection of AAVrg-Con/Fon-ChR2-EYFP into the SNr/SNc (blue). Roman numerals and colored boxes indicate regions of interest displayed in **F**. N = 3 mice.

**F**, A unilateral injection of AAVrg-Con/Fon-ChR2-EYFP into the SNr/SNc of Vglut2+/Vgat+ mice results in the detection of cell bodies (indicated by white arrows) in the STN (I; brown) and the GPi (II; red). N = 3 mice.

Abbreviations: GPi: globus pallidus interna, LHb: Lateral habenula, STN: subthalamic nucleus, int: internal capsule, cpd: cerebal peduncle, SNc: Substantia nigra pars compacta, SNr: Substantia nigra pars reticulata, ML: mediolateral, AP: anteroposterior.

**Supplementary Figure 5 | Sst+ GPi-LHb neurons send collaterals to both GPe and SNc.**

**A**, Upper left: Schematic representation of the experimental strategy: AAVrg-Cre virus injected in the Lhb, AAVrg-DIO-EGFP virus injected in the SNc and AAVrg-DIO-mCherry virus injected in the GPe of wt animals. Lower right: Representative ISH images showing the double labeled Sst+ cells in the GPi. Right column: Close up images of the double labeled neurons in the GPi. N = 3 mice.

**B**, Representative images showing the double-labeled terminals in the LHb.

**C**, Upper left: Schematic representation of the experimental strategy: AAVrg-DIO-TetLCh-dTomato virus injected in the LHb, AAVrg-CreOn-FlpOn-EGFP virus injected in the SNc and

AAVrg-DIO-flpo virus injected in the GPe of Sst-cre animals. Lower right: Representative images showing the double labeled cells in the GPi. N = 3 mice.

**D**, Close up images of the double labeled neurons in the GPi.

**E**, Upper: Representative image of EGFP-labeled terminals in LHb. Lower: Representative image of EGFP-labeled terminals in GPe.

Abbreviations: wt: wild type, MHb: Medial habenula, LHb: Lateral habenula, GPi: globus pallidus interna, SNc: Substantia nigra pars compacta, CPu: Caudate putamen.

**Supplementary Figure 6 | Electrophysiological and immunohistological characterization of Vglut2+/Vgat+ GPi neurons input to the CPu and the GPe.**

**A**, ISH images of the rabies tracing experiment in A2A-expressing CPu neurons. No Rabies-expressing monosynaptic input neurons could be detected in the GPi. Related to **Figure 2B**. Scale bar: 100μm.

**B**, ISH images of the rabies tracing experiment of Oprm1-expressing CPu neurons. Left: Representative Rabies-expressing neuron (green) in the GPi targeting Ormp1-expressing CPu neurons co-labeled with SST (red) by immunohistological staining. Right: closeup of the neuron indicated to the left. Related to **Figure 2C**. Scale bar: 5μm.

**C**, Scatter plot showing the correlation between the jitter of GABA and AMPA currents, demonstrating a high Pearson correlation coefficient (r = 0.794). Each point represents an individual recording, with jitter values plotted for both neurotransmitter currents.

**D**, ISH images identifying patched and biocytin-filled (turquoise; n = 9), neurons in the GPe showing expression of PV (pink) and not FoxP2 (green). Scale bar: 20μm.

**E**, Rheobasic bursting response (left) of a PV-expressing GPe neuron and two-times rheobasic response (right) of another PV-expressing GPe fast spiking neuron.

**F**, Spontaneous burst oscillation of PV-positive GPe neuron upon hyperpolarization to -90 mV.

**G**, Illustration depicting the experimental strategy: AAV-Con/Fon-GCaMP6m was unilaterally injected directly into the GPi of Vglut2-Cre/Vgat-Flp mice.

**H**, Up: Representative images of immunostaining of Vglut2 (green) and Vgat (red) on GCaMP6m-expressing GPi neuron terminals (white) in the main projection targets regions of Vglut2+/Vgat+ GPi-LHb neurons (from left to right: CPu (light blue), GPe (blue), LHb (pink), and PFC (green)). Scale bar: 10μm. Down: close-ups of area indicated in the top images, displaying the overlay of Vgat, Vglut2 and GCaMP6m-expressing GPi efferents in the Z-plane. Left, down; Identified bouton in the coronal plane, the white arrow indicates the area of cross-sectioning in the z-plane displayed to the right. Right, down; The imaged z-plane of the bouton indicated to the left, for each imaged channel (white: GCaMP6m anti-GFP; green: Vglut2; red: Vgat). Scale bar: 1µm.

**I**, Proportion of GCaMP6m-expressing GPi boutons co-expressing Vglut2 and Vgat within the four main projection targets of Vglut2+/Vgat+$^{GPi\text{-}LHb}$ neurons (proportion of total boutons identified by the expression of the viral fluorophore: light blue; CPu (n = 65/75), blue; GPe (n= 274/292), pink; LHb (n = 433/465), green; PFC (n = 31/34)). Boxplot: center lines, median; edges, upper and lower quartiles; whiskers, largest and smallest value no more than 1.5×IQR of the edges. Each dot represents the proportion of GPi boutons co-expressing Vglut2 and Vgat for each analyzed brain section, color-coded by the brain area analyzed.

Abbreviations: Lhb: Lateral habenula, GPi: globus pallidus interna, CPu: Caudate putamen, PFC: prefrontal cortex.

**Supplementary Figure 7 | Fiber placement and virus expression in behavioral cohorts.**

**A**, Left: Schematic representation of the experimental strategy: Sst-cre mice were injected with AAVrg-DIO-jGCaMP7s in LHb and implanted with optical fibers in GPi. Right: Images depicting fiber placement and jGCaMP7s expression (green) in individual mice.

**B**, Left: Schematic representation of the experimental strategy: Pv-cre mice were injected with AAV-DIO-jGCaMP7s and implanted with optical fibers in GPi. Right: Images depicting fiber placement and jGCaMP7s expression (green) in individual mice.

**C**, Left: Schematic representation of the experimental strategy: Sst-cre mice were injected with AAVrg-DIO-TeTxLC-dTomato in the LHb. Right: Images from a representative mouse depicting TeTxLC-dTomato-expressing neurons (red) in the GPi.

Abbreviations: GPI: Globus pallidus interna, ic: internal capsule.

**Supplementary Figure 8 | Sst+ and Pv+ GPi populations are not responding to negative values.**

**A**, Auditory fear conditioning of Pv-Cre (top) and Sst-Cre (bottom) mice. Traces (left) show the fluorescence response (blue, red, and magenta) and the probability of mice being stationary (black) for trials of the second habituation, the conditioning, and the extinction sessions (left to right, see event pictograms). Mice were considered stationary when median body and head velocity in a 300 ms window averaged lower than 1 cm/s. Grey shading: auditory conditioned stimulus (CS) presentation. Mean ± SEM, traces pooled over animals (N = 3 per group). Regressions (right) assess the relation of fluorescence and immobility during the CS period. Circles: single-trial CS responses, trial type color-coded following the traces. Line and shading:

mean ± SEM of the regression fits. *: $p<0.001$, Wald test of the main effect of the slope; linear mixed-effects model. N = 3 mice per group.

**Supplementary Figure 9 | Sst+ GPi-LHb neuronal activity does not encode valence or prediction error signaling in the single-reversal maze task.**

**A**, Schematic illustration of our setup, showing the regions where we track the changes in the activity patterns following rewarded and non-rewarded trial outcomes pre- vs post-reversal of the reward location.

**B**, Sst+ ΔF following rewarded and non-rewarded trial outcomes (time = 0, dashed line) on day 16, the second day post-reversal. Baseline for ΔF: F at the moment of the outcome delivery. Inset: angular velocity in the same time frame. Traces: mean ± SEM. Vertical lines: median times at which mice came to a halt at the spout. Gray shading: 1s time window used to compute means in panel C. Rewarded/non-rewarded n = 122/85 trials (pooled, N = 5 mice).

**C**, Mean ΔF in the second after outcome delivery, split by outcome and plotted across days. Thin lines: individual Sst-Cre mice. Circles: mean ± SEM. No significant differences between positive and negative outcome responses detected (all bins $p > 0.1$; paired t-tests, no correction). N = 5 mice. Inset: scatterplot and regression line depicting the relationship between the mean angular velocity and the mean ΔF in the 1s window following non-rewarded trial outcomes. Points: session means. n = 81 sessions (pooled, N = 5 mice). Note that there are sessions missing photometry data and 100%-accurate sessions without non-reward outcomes.

**D-E**, same as B-C for Pv-Cre mice.

**D**, Rewarded/non-rewarded n = 103/39 trials (pooled, N = 3 mice).

**E**, All bins $p > 0.1$; paired t-tests, no correction. N = 3 mice. Inset: n = 52 sessions (pooled, N = 3 mice).

**F**, Motif usage aligned on the end of the motif 41 (red). Motif 41 is frequently followed by motif 5 (in blue) in the pre-reversal phase.

**G**, Fraction of trials completed plotted against session time for training day 14. Colorbar on y-axis: trial quartile bins (Q1-4). Thin lines: individual mice. Thick line: mean. Inset: total number of trials in session. Circles: individual mice. Bar: mean ± SEM. N = 8 mice (5 Sst-Cre + 3 Pv-Cre).

**H**, Left: Mean trial rates in each trial quartile bin (background shading, colors as in **G**). Right: Mean accuracy in each trial quartile bin. Thin lines: individual mice. Circles: mean ± SEM. *: $p < 0.001$; repeated-measures ANOVA. N = 8 mice.

**I**, Exemplary session snippet showing Sst+ GPi-LHb population activity, and the mouse’s total and angular velocity over the course of several trials, post-reversal (day 15). Background shading:

inferred behavior (see legend). Dashed lines: drop icons indicate trial outcomes (rewarded or non-rewarded) and arrowheads entries into the maze junction (up-arrow: choice run, down-arrow: initiation run). Data from one Sst-Cre mouse.

**Supplementary Figure 10 | Sst+ GPi-LHb neuronal activity aligned to motif onset in the single-reversal task.**

**A**, Line graphs reporting the mean ± SEM of calcium transients signals recorded through fiber photometry and aligned to motifs onset during the pre- (in blue) and post-reversal (in orange) phase.

**Supplementary Figure 11 | Supplementary behavioral data illustrating the behavioral changes following the silencing of Sst+ GPi-LHb neurons during the single-reversal task.**

**A**, Visual explication of rewarded and non-rewarded transitions.

**B**, Time immobile, **C** Number of total transitions, **D** Number of rewarded transitions, **E** Number of errors of AAVrg-DIO-dTomato and AAVrg-DIO-TeTxLC-dTomato injected mice in the single-reversal task across 20 days of test. Mean ± SEM. *: $p < 0.05$, multiple comparisons using the Mann-Whitney U test followed by Holm-Sidak correction.

**Supplementary Figure 12 | No dynamic activity modulation was detected in GPi Pv+ neurons during the single-reversal task.**

**A**, Raster: GPi Pv+ population activity (F) for individual pre- and post-reversal choice runs. Trials were grouped by task stage and turn direction, sorted by fraction of prior trials completed in their session, and finally binned into quartiles (Q1-4). The bins are unevenly sized due to the exclusion of trials in which mice returned to the center corridor before reaching a reward zone. Data from sessions 1-4 and pre-reversal ipsiversive trials omitted, the latter due to the low number of trials available. Traces: mean ± SEM for each raster bin. n = 841/151/506 trials (pooled, N = 3 mice).

**B**, Top row: stacked histograms indicating the fraction of contra- vs ipsiversive choices, by session quarter, over the course of the experiment. Quarters are based on trial quartiles, not time. Trials pooled over Pv-Cre mice. Middle and bottom rows: density plots visualizing Pv+ activity (F) during contra- (middle) and ipsiversive (bottom) choice turns, by session quarter, over the course of the experiment (same as above). One data point, mean F, included per turn. Circles: mean ± SEM of turns by session quarter; mean omitted for bins with fewer than 5 data points total. Right-hand histograms: distribution of mean turn F pooled over all sessions. Contra/ipsi n = 1202/620 trials (pooled, N = 3 mice; cf. Figure 6G).

**C**, Top: probability of ipsiversive choices plotted against the average F during contraversive turns for all session-quartile bins. Small circles: individual session-quartile averages; contra turn F z-scored per mouse. Black circles: median and IQR of binned session-quartiles; bin edges as indicated in bottom panel. n = 196 session-quartiles (pooled, N = 3 mice). Bottom: same as above, plotted per mouse. Thin lines: median of binned session-quartiles for individual mice; bin edges are indicated on the x axis. Bars: mean ± SEM. $p = 0.6$, repeated-measures ANOVA. N = 3 mice.

### METHODS

**Viral Constructs.** *Anatomy and histology*.
AAVrg-DIO-ChR2-mCherry; pAAV-Ef1a-double floxed-hChR2(H134)-mCherry, Addgene, cat. no. 20297 (7x10e12 viral genomes/ml).
AAVrg-Con/Fon-ChR2-EYFP; pAAV-hSyn-Con/Fon-hChR2(H134)-EYFP, Addgene, cat. no. 55645 (7x10e12 viral genomes/ml).
AAVrg-FLEX-jGCaMP7s; pGP-AAV-syn-FLEX-jGCaMP7s-WPRE, Addgene, cat. no. 104491 (7x10e12 viral genomes/ml).
AAV8-Con/Fon-ChR2-EYFP; pAAV-hSyn-Con/Fon-hChR2(H134)-EYFP, Addgene, cat. no. 55645 (2x10e13 viral genomes/ml).
AAV8-Con/Fon-GCaMP6m; pAAV-EF1a-Con/Fon-GCaMP6m, produced in the laboratory of Dr. Karl Deisseroth (Stanford University), AAV#1746 (3,5x10e12 viral genomes/ml).
AAVrg-Cre; pAAV-EF1a-Cre, Addgene, cat no 55636 (2.1x10^13 viral genomes/ml).
AAVrg-FLEX-EGFP; pCAG-FLEX-EGFP-WPRE, Addgene, cat no 51502 (2.3x10e13 viral genomes/ml).
AAVrg-DIO-mCherry; pAAV-EF1a-double floxed-hChR2(H134)-mCherry, Addgene, cat no 20297 (1.2x10^13 viral genomes/ml).
AAVrg-CreOn-FlpOn-EGFP; pAAV-hSyn-Con/Fon hChR(H134)-EYFP, Addgene, cat no 55645 (1.2x10e13 viral genomes/ml).
AAVrg-DIO-flpo; pAAV-EF1a-Flpo, Addgene, cat no 55637 (2,2x10^13 viral genomes/ml).

*Rabies tracing*.
The rabies helper virus AAV5-DIO-TVA-V5-RG; pAAV-EF1a-DIO-TVA-V5-t2A-RG, was created as described previously[46] (Addgene cat. plasmid no. 119743). AAV5 viral particles were produced by the Virus Vector Core facility at the University of North Carolina (1.2x10e13 viral particles/ml). The EnvA pseudotyped Rabies Glycoprotein-deleted Rabies Virus (Rb) (1x10e10 infection unit/ml) was produced in-house as described previously[46].

*Tetanus Toxin.*
AAVrg-FLEX-TeTxLC-dTomato; pAAV-hSyn-FLEX-TeLC-P2A-dTomato was a gift from Sandeep Datta, Addgene, cat. plasmid no.159102; AAVrg was produced by the Virus Vector Facility at the University of Zurich (7x10e12 viral particles/ml).

*GCaMP expressing*.

AAVrg-FLEX-jGCaMP7s; pGP-AAV-syn-FLEX-jGCaMP7s-WPRE, Addgene, cat.no. 104491 (7x10e12 viral genomes/ml).

AAV1-FLEX-jGCaMP7s; pGP-AAV-syn-FLEX-jGCaMP7s-WPRE, Addgene, cat. no. 104491 (1x10e13 viral genomes/ml).

**Animals.** Experiments were conducted using adult male and female mice, wild type C57BL/6J (Charles River Laboratories) or transgenic mouse lines; Vglut2-Cre: $Slc17^{a6tm2(cre)Lowl}$, Jackson stock no. 028863; Sst-Cre: $B6N.Cg\text{-}Sst^{tm2.1(cre)Zjh}$, Jackson stock no. 018973, Pv-Cre: $Pvalb^{tm1(cre)Arbr}$, Jackson stock no. 008069. Vgat-Flpo transgenic mice were produced by Hongkui Zeng (Allen Institute for Brain Science, USA).

All transgenic mice used in experiments were heterozygous for the transgenes. Mice were maintained under standard housing conditions with a 12-hour light cycle and with *ad libitum* access to food and water unless placed on a food or water restriction schedule (see behavioral paradigms for details). All procedures were approved and performed in accordance and compliance with the guidelines of the Stockholm Municipal Committee (approval no. N166/15, 7362-2019 and 15440-2020).

Animal cohorts

*Anatomy and histology.*: N=7 retro LHb injection Vgat-Flpo/Vglut2-Cre, N=3 retro LHb injection retro-AAV-DIO-mChery in Sst-cre, N=3 retro SNr/SNc injection Vgat-Flpo/Vglut2-Cre, N=6 direct injection GPi Vgat-Flpo/Vglut2-Cre, N=6 retro LHb injection of GCaMP7s in Sst-Cre mice, N=3 retro viral strategy of Supplementary Figure 5 in wt, N=3 retro viral strategy of Supplementary Figure 5 in Sst-Cre.

*Electrophysiology*: N=8 Vgat-Flpo/Vglut2-Cre mice.

*Rabies*: N=3 Rabies injection in the CPu of A2A-Cre, N=6 Rabies injection in the CPu of Oprm1-Cre, N=3 Rabies injection in the SNc of DAT-Cre mice.

*Fiber photometry*: N=10 Sst-Cre mice and N=6 Pv-Cre mice.

*Tetanus Toxin:* N=16 Sst-Cre mice (controls) and N=15 Sst-Cre mice (TeTxLC).

See also **Supplementary Table 2**.

Animals were excluded from further analysis if histological inspection revealed nonspecific viral expression or incorrectly positioned fiber implants. Specifically, animals were removed from the study if: (i) viral expression was observed outside the intended target region (e.g., in adjacent nuclei or non-target cell populations); (ii) expression within the target region was sparse, incomplete, or uneven; (iii) fiber optic implants were misplaced and did not overlay the labelled neuronal population.

**Viral injections and implants.** *General procedure***.** Mice were anesthetized with isoflurane (2%) and placed into a stereotaxic frame (Harvard Apparatus, Holliston, MA). Before the first incision the analgesic Buprenorphine (0.1 mg/kg) and local analgesic Xylocain/Lidocain (4 mg/kg) were administered subcutaneously. The temperature of the mice was maintained at 36 °C with a feedback-controlled heating pad. For viral injections a micropipette attached on a Quintessential Stereotaxic Injector (Stoelting, Wood Dale, IL) was used. The pipette was held in place for 5 min after the injection before being slowly retracted from the brain. The analgesics Carpofen (5mg/kg) was given at the end of the surgery, followed by a second dose 18-24h after the surgery.

*Anatomy*. For anatomical characterization of the GPi-LHb pathway in Sst-Cre mice, 0.3µl of either AAVrg-DIO-ChR2-mCherry or AAVrg-DIO-jGCaMP7s was unilaterally injected into the LHb (coordinates: AP -1.65 mm, ML 0.92 mm 10◦, DV -2.45 mm). Targeting of the GPi-LHb pathway in Vgat-Flp/Vglut2-Cre mice was achieved by either an unilateral injection of 0.3µl of AAVrg-Con/Fon-ChR2-EYFP into the LHb (coordinates: AP -1.65 mm, ML 0.92 mm 10◦, V -2.45 mm) or a direct injection of 0.2µl of AAV8-Con/Fon-ChR2-EYFP or AAV8-Con/Fon-GCaMP6m in the GPi (coordinates: AP -1.22 mm, ML 1.75 mm, DV -4.1 mm). Targeting the GPi-LHb neurons that project to GPe and SNc in wt animals was achieved by injection of AAVrg-Cre in LHb (coordinates: AP -1.65 mm, ML 0.92 mm (10°), DV -2.45 mm), AAVrg-DIO-mCherry in GPe (coordinates: AP -0.22 mm, ML 1.75 mm, DV -3.35 mm) and AAVrg-DIO-EGFP in SNc (coordinates: AP -3.08 mm, ML 1.25 mm, DV -4.25 mm). Labelling the Sst+ GPi-LHb neurons that project to GPe and SNc in Sst-cre animals was achieved by injection of AAVrg-DIO-TetLCh-dTomato in LHb (coordinates: AP -1.65 mm, ML 0.92 mm (10°), DV -2.45 mm), AAVrg-DIO-flpo in GPe (coordinates: AP -0.22 mm, ML 1.75 mm, DV -3.35 mm) and AAVrg-CreOn-FlpOn-EGFP in SNc (coordinates: AP -3.08 mm, ML 1.25 mm, DV -4.25 mm).

*Rabies tracing.* For cell-type specific monosynaptic retrograde tracing mice were unilaterally injected with 0.3µl of the helper virus AAV-DIO-TVA-V5 in the SNc (DAT-Cre; coordinates: AP -3.08 mm, L 1.2 mm, V- 4.25 mm) or 0.4µl in the CPu (A2A-Cre or Oprm1-Cre; coordinates: AP +0.5 mm, L 2.1 mm, V- 2.0 mm). After 21 days, 0.3µl of Rabies-EGFP virus was injected at the same coordinates. Mice were perfused and brains were collected for histology 7 days post rabies injection. To ensure that pseudotyped rabies virus infection is strictly dependent on prior helper virus expression, we performed control experiments in C57BL/6 mice, which showed negligible infection in the absence of helper virus (see **Supplementary Information** for details).

*Slice electrophysiology.* Targeting and labeling of neuronal inputs was achieved by a unilateral injection of 0.2µl AAV8-Con/Fon-ChR2-mCherry into the GPi (coordinates: AP -1.22 mm, ML 1.75 mm, DV -4.1 mm) of Vgat-Flp/Vglut2-Cre mice.

*Fiber Photometry.* Adult male and female Sst-Cre mice were injected with 0.3µl of AAVrg-DIO-jGCaMP7s[21] into the LHb (coordinates: AP -1.65 mm, ML ±0.95 mm (10°), DV -2.45 mm). PV-Cre mice were injected with 0.2µl of AAV1-DIO-jGCaMP7s directly into the GPi (coordinates: AP -1.22 mm, ML ±1.75 mm, DV -4.1mm). Injected mice were implanted one week later with optical fibers (200µm diameter) aimed directly above the GPi (coordinates: AP -1.22 mm, ML ±1.75 mm, DV -3.9 mm).

*Tetanus toxin.* Male and female Sst-Cre mice were bilaterally injected with 0.3µl of AAVrg-DIO-TeTxLC-dTomato[24] in the LHb (coordinates: AP -1.65 mm, ML ±0.95 mm (10°), DV -2.45 mm). Training of the mice in the single-reversal maze task started at least 21 days post injection.

**Histology.** *General procedure***.** Mice were deeply anesthetized with pentobarbital and then transcardially perfused with 0.1M PBS followed by 4% paraformaldehyde in PBS 0.1M. Brains were removed and post-fixed in 4% paraformaldehyde in PBS 0.1M overnight at 4°C and then washed and stored in 0.1M PBS. Coronal (60µm thickness) or sagittal sections (80µm thickness) were cut using a vibratome (Leica VT1000, Leica Microsystems, Nussloch GmbH, Germany).

*Immunostaining***.** Immunostaining was performed on free-floating sections. Briefly, sections were incubated for one hour in 0.3% TritonX-100 in Tris-buffered saline (0.3% TBST). For difficult stainings, the brain sections were treated with a preheated (40°C) antigen retrieval solution (10mM sodium citrate, 0,05% Tween20, pH:6) for 1-2 minutes. To block non-specific antibody binding, sections were incubated in 5% Normal Donkey Serum in 0.3% TBST, for one hour at room temperature (RT). Sections were subsequently incubated overnight at RT with primary antibodies followed by a 4h incubation with the secondary antibodies. The sections where thereafter washed consecutively in 0.3% TBST, 1x TBS and 1x PBS (15min each) and mounted on glass slides (Superfrost Plus, Thermo Scientific) and coverslipped (Thermo Scientific) using glycerol: 1x PBS (50:50).

*Primary antibodies used* (*abbreviation; dilution; company; catalogue number*): Guinea pig anti-parvalbumin (PV; 1:1000; Synaptic Systems; 195 004), Goat anti-somatostatin (SST; 1:500; Santa Cruz Biotechnology; sc-7819), Rabbit anti-FoxP2 (FoxP2, 1:1000; Abcam; ab16046), Chicken anti-V5 (V5, 1:1000, Novus Biologicals, NB600-379), Guinea pig anti-VGluT2 ( VGLUT2, 1:1000, Synaptic Systems, 135 404), Rabbit anti-VGAT (VGAT, 1:1000, Synaptic Systems, 131 002), Rabbit anti-Mu Opioid Receptor (MOR; 1:200; Abcam; ab134054), Sheep anti-Tyrosine Hydroxylase (TH, 1:1000; Millipore; AB1542), Rabbit anti-Green Fluorescent Protein (GFP; 1:1000; Invitrogen; A6455), Chicken anti-Green Fluorescent Protein (GFP; 1:1000; Invitrogen; A10262), Goat anti-Green Fluorescent Protein (GFP; 1:1000 Abcam; ab5450), Rabbit anti-Red fluorescent Protein (RFP; 1:1000; Rockland; 600401379).

*Secondary antibodies used* (*fluorophore; dilution; company; catalogue number*): Donkey anti-rabbit (Cy3; 1:1000; Jackson; 711-165-152), Donkey anti-rabbit (Cy5; 1:1000; Jackson; 711-175-152), Donkey anti-chicken (Cy3; 1:1000; Jackson; 703-165-155), Donkey anti-chicken (Cy5; 1:1000; Jackson; 703-175-155), Donkey anti-guinea pig (Alexa Fluor 405; 1:500; Jackson; 705-475-147), Donkey anti-guinea pig (Cy3; 1:1000; Jackson; 706-165-148), Donkey anti-guinea pig (Cy5; 1:1000; Jackson; 706-175-148), Donkey anti-goat (Alexa Fluor 488; 1:1000, Jackson; 705-545-147).

**Anatomical and histological analysis.** *Anatomical mapping of GPi-LHb axons and cell bodies.* For viral constructs and strategies see *Viral Constructs*. Tiled whole brain images of sagittal or coronal cut sections were acquired at 10x magnification and close-up z-stacked images were acquired at 20-63x magnification using a Zeiss (LSM800) confocal microscope.
For the whole-brain axonal patterning plots, each sagittal cut brain section was manually mapped to a ML coordinate and x-y coordinate for each fluorescent image pixel (axonal fluorescent labeling) was segmented out in ImageJ. Cell bodies were manually mapped using the CellCounter in ImageJ. A 2D reconstruction of the axonal arborization/cell bodies was thereafter plotted within the Allen Reference Atlas framework in R (**Supplementary Figure 1** and **Supplementary Figure 4**). For axon location confirmation within a target region, immunohistological labeling was performed on coronally sectioned brains (**Figure 1**)**.**

*Quantification of signal intensity*. Tiled whole brain coronal images were acquired for each animal using a Leica epifluorescence microscope. Images were down sampled and aligned to the Allen

Reference Atlas using cell registration software in Napari, developed in-house. Regions of interest were designed in the target areas and images were inverted and thresholded on ImageJ. Heatmaps were created after normalization of the values to the background on ImageJ and superimposed on Allen Reference Atlas (**Figure 1J-M** and **Supplementary Figure 3**).

*Neurotransmitter labeling in axonal terminals***.** Immunostaining was performed to quantify the co-labeling of GPi boutons with Vgat and Vglut2. GPi boutons were identified by injecting AAV-Con/Fon-GCaMP6m directly into the GPi of Vglut2-Cre/Vgat-Flp mice. 1-3 Z-stack images were captured of each brain region of interest on a Zeiss (LSM800) confocal microscope. Images were acquired using identical pinhole, gain, and laser settings for all brain regions and analyzed using ImageJ and/or Imaris software. Boutons were identified by GCaMP6m expression and the fluorescence intensity of the Vglut2 and Vgat staining within the area of each identified bouton was measured.

*Fiber placement and virus expression in behavioral cohorts.* The brains of mice in the fiber photometry and tetanus toxin cohorts were collected as specified in *General procedure*, cut (60µm thickness) coronally and all consecutive sections covering the GPi and adjacent brain regions were collected and mounted. Assessment of fiber placement was based on the lesion from the fiber in the tissue. Virus expression level and specificity was manually evaluated and animals with misplaced fibers or lack of virus expression/specificity were excluded from the study.

**Behavior.** Behavioral experiments were performed in a consistent sequence to minimize potential carryover effects. Mice first underwent the nose-poke serial reversal task, followed by the single-reversal maze task, then the open field test, and finally, only a subset of fiber photometry animals participated in fear conditioning (see **Supplementary Table 2**). All Sst-Cre dTomato and Sst-Cre TeTxLC animals completed the single-reversal maze task before the exploration of the open field arena

*Fiber photometry, pose estimation and motifs segmentation.* The fiber photometry calcium imaging setup was as previously reported[47]. We acquired calcium-dependent fluorescence, elicited at jGCaMP7's[21] optimal excitation wavelength (470 nm) and indicative of neuronal activity, as well as interleaved calcium-independent control signals at the calcium indicator's isosbestic wavelength (405 nm). The light power of the 470 nm LED was dialed to about 20-30 µW, with the power of the 405 nm LED ranging slightly lower, usually between 10-25 µW.

Alongside the photometry recordings, we captured videos of the mice's behavior from below at 60 frames per second. Using DeepLabCut software[22,48] for post-hoc analysis, we extracted the xy-coordinates of three key body parts (snout, center of gravity, and tail base) and, in a separate analysis, of six body parts (snout, four paws, and tail base). For alignment purposes, video and photometry frames, as well as important task events, were timestamped, usually using the Bonsai software[49], which communicated with external hardware via an Arduino Micro microcontroller (Arduino) running the firmata protocol[50]. The photometry traces and the behavior videos were recorded at varying frame rates (see below). Before further processing, we therefore uniformly downsampled all photometry and locomotor tracking data recorded at higher frame rates to the 10 Hz rate of most of the photometry recordings, by binning and averaging.

Each photometry recording was further preprocessed as follows: The 405 nm control signal was first scaled to the 470 nm neuronal activity signal, using linear least-squares regression, and then subtracted from it. This removed motion and autofluorescence-related artifacts. Next, we estimated the baseline fluorescence of the raw activity signal over time, by fitting a least-squares regression line to the scaled control signal and then divided the corrected activity signal at each time point by the corresponding raw baseline estimate. This resulted in a normalized and photobleaching-adjusted, motion and autofluorescence-corrected neuronal activity trace ($\Delta F/F$). Finally, we computed the rolling z-score (5 min moving window) of this trace to remove very slow activity transients, as well as to standardize the measurements across animals (F in our figures).

The downsampled spatial trajectories of all tracked body parts were further preprocessed using a Kalman smoother, assuming constant movement velocity frame-to-frame. Based on the smoothed trajectories, we computed the velocity of the body mid-point (cm/s) as well as its angular velocity (°/s) relative to the base of the tail. To segment out left and right turns, forward motion and pauses, we used our published optimization algorithm[25]. For the segmentation of the behavior in motifs, the unsupervised deep learning framework VAME[23] was trained on 51 videos for the open field test and 76 videos for the single-reversal maze task with a test fraction of 0.1, a time window of 30 frames, and a latent dimensionality of 30, converging at 50 epochs. Clustering analysis with 100 discrete states and a 1% motif usage threshold identified 45 (open field test) and 49 (single-reversal maze task) relevant behavioral motifs within the dataset.

*Nose-poke serial reversal task.* Food-restricted mice, kept above 85% of their free-feeding body weight, were trained on an established probabilistic serial reversal task[13,16,25,26]. In brief, mice initiated trials at a central nose poke port, then chose between two identical ports located to its left and right. A nose poke into the “correct” port resulted in a liquid sucrose reward (15% sucrose,

3.75 µl) with 75% probability, whereas the “incorrect” port yielded nothing. The roles assigned to the choice ports reversed every 7-23 rewarded trials, at random and without indication. LED lights mounted inside the ports signaled only whether the initiation or choice ports were actionable. To facilitate learning of the full task, we pre-trained mice on a version of the task where uninitiated pokes of the correct choice port, as well as initiation port pokes, yielded water rewards. In this simplified paradigm, both the initiation and the correct choice port were lit, whereas the incorrect choice port was not, to unambigiously signal the location of the reward. Mice were transitioned to the full task once they consistently nose poked for rewards throughout the session. Photometry recordings commenced once the mice reached asymptotic, above chance performance. Nose pokes were recorded as infrared beam breaks by a pyboard microcontroller (pyboard.org), which operated the 15x15 cm custom-built operant chamber. The pyboard also registered every recorded photometry and video frame. Training and recording sessions lasted 30 to 60 minutes. A BlackFly USB3 camera (Flir) captured video from above at 20 or 40 Hz, whilst fiber photometry fluorescence data was acquired at half these rates.

We fitted relative action values using Q-learning, as reported previously[25]. The relative action value was computed as $\beta \times (Q_{contra} - Q_{ipsi}) + b$, where $Q_{contra}$ and $Q_{ipsi}$, $\beta$, and $b$ denote the Q-values of the contralateral and ipsilateral ports, the explore-exploit trade-off, and the static port bias, respectively. The relative action values are the log-odds of the animal choosing the ipsilateral port, such that negative values favor ipsilateral port and positive values favor contralateral port choices. Reward prediction errors were calculated as the relative action value shift ($\Delta Q$) in favor of the action taken on each trial. Importantly, a relative action value decrease was treated as a positive error, and a relative action value increase was treated as a negative error, on leftward but not rightward trials.

In **Figures 4K** and **R**, we fit a number of Ridge regressions to assess which variable best explained mean F in the second following choice port entries – $\Delta Q$, outcome, or the fraction of time spent nose-poking (labeled “% in port”). To obtain cross-validated variance explained scores ($cvR^2$) and to estimate optimal regularization penalties, we used 10-fold cross-validation throughout. We used a published variance analysis[51] approach to parcel out the maximum and unique explanatory power of each variable. Firstly, to compute a maximum $cvR^2$ for each variable individually, we fit regressions in which all variables except for the variable of interest were shuffled. Secondly, to estimate how much $cvR^2$ is uniquely explained by each variable, we fit regressions in which only the variable of interest was shuffled and computed the loss of power ($\Delta R^2$) vis-a-vis a fully unscrambled model (labeled “combined”).

*Auditory fear conditioning.* The conditioned stimulus (CS) was a 10 s, 4kHz, pulsed tone (250 ms on/off) measuring 55 dBA inside the chamber; the unconditioned stimulus (US) was a 1 s, 0.3 mA foot shock. Each day, mice were exposed to 20 presentations of the CS, separated by random inter-trial-intervals (10-70 s). On days 1 and 2 (habituation), no US was delivered. On day 3 (conditioning), every CS was followed by the US. On day 4 (extinction), a plastic mat covered the grid floor. In this novel environment, the CS was once again presented without the US.

The task was controlled by an Arduino Micro microcontroller (Arduino) equipped with a piezo buzzer. The shocks were generated by a TTL-triggered aversive stimulator (ENV-414S, Med Associates) connected to the grid floor of the conditioning chamber (ENV-307W-CT, Med Associates). The chamber was located inside an unilluminated, sound-attenuating cubicle (ENV-016MD, Med Associates). Videos were recorded from above at 30 Hz, using an infrared webcam. Fiber photometry fluorescence data was acquired at 10 or 20 Hz.

*Open field.* Overnight food or water-deprived mice were left to freely explore a 49x49 cm box with a clear plastic floor for 15 to 30 minutes. Except for red and infrared light sources, all lights in the room were dimmed. A BlackFly USB3 or an Oryx 10GigE camera (both Flir) recorded video from below at 60 Hz. Fiber photometry fluorescence data was acquired at 10 Hz.

*Single-reversal maze task.* The single-reversal maze task consisted of a 40x40 cm box with a clear plastic floor and a number of removable inserts. To fashion 5 cm-wide corridors along the 4 walls of the box and a diagonal corridor crossing the center, we inserted two “half pyramid”-shaped (triangular base, 2 sides slanted) inserts of clear plastic into the box. The slanting of the inserts’ walls towards the center of the box allowed the photometry patch cable to move freely without catching on corners or edges. Inserts equipped with water spouts were placed in three corners of the box, blocking off a corner at one end of the center-crossing corridor and the two corners next to it. The open fourth corner, together with the three adjoining corridors, created an arrow-shaped maze, resembling a pointed T-maze. Various visual cues, visible through the clear plastic throughout the arena, aided navigation. Facing the 3-way corner, the box’s walls left of the center corridor were white, the walls on the right black. Moreover, each wall featured additional visual cues of different shapes and colors and a DUPLO panda figure (Lego) was placed at the center of the left half pyramid insert. The whole arena was illuminated by a white LED light strip glued along the rim of the clear plastic floor.

To motivate foraging for water rewards, mice were water-restricted, receiving 1 ml of water per day in total (in-task and researcher-supplied). Every 12 days of water restriction were followed by 2 days of rest with ad-libitum water access.

Water-restricted mice were first habituated to the maze for 3 days. During habituation we removed the spout-equipped inserts, thus making all corridors and corners accessible. In the following 14 experimental days (i.e. excluding rest days), mice were trained to shuttle back and forth between the spouts at the ends of the center “initiation” corridor and one of the “choice” corridors, where water rewards (13 µl) were delivered upon approach in alternating fashion. Approaching the spout in the “incorrect” choice corridor reset the reward location to the initiation spout without yielding a reward. Approaching the same spout repeatedly, moving between choice spouts, or entering corridors without approaching the spout at the end did not advance the task. Consequently, a “choice trial” ends once the first choice spout is reached after trial initiation, whereas a “trial initiation” ends once the initiation spout is reached after a choice trial. Choice accuracy (**Figure 5C**) was calculated as the number of correct choice trials divided by the total number of choice trials. Task efficiency was calculated by considering all transitions between spouts, including those that do not advance the task (**Supplementary Figure 11A**): number of rewarded transitions/(number of rewarded transitions + number of non-rewarded transitions). For the fiber-implanted mice, the correct choice corridor was on the side contralateral to their implant, whereas for the tetanus toxin-expressing mice, the correct choice was the right corridor, at this stage. The two choice corridors swapped roles, the rewards now being obtainable in the formerly incorrect arm, in the subsequent reversal phase of 6 days. Other than the rewards, no cues signaled the correct choice or the reversal. All sessions lasted 20 minutes and for fiber-implanted mice the neuronal activity was recorded during all sessions (from day 1 to day 20).

The task was implemented using the Bonsai software[49], used to detect entries into the corridor end zones from a live video feed, and two Arduino Micro microcontrollers (Arduino), controlling the water spouts and communicating with Bonsai via the firmata protocol[50], respectively. Video frames were acquired at 60 Hz using an Oryx 10GigE camera (Flir) placed below the arena. Fiber photometry fluorescence data was acquired at 10 Hz.

**Electrophysiology.** Vglut2-Cre/Vgat-Flpo mice were injected with AAV-Con/Fon-ChR2-EYFP at 8-9 weeks and recorded 10-12 weeks of age. 250 µm thick coronal slices were cut with a vibratome (VT1200S, Leica, Germany) in ice-cold cutting solution, containing (in mM): 40 NaCl, 2.5 KCl, 1.25 NaH2PO4, 26 NaHCO3, 20 glucose, 37.5 sucrose, 20 HEPES, 46.5 NMDG, 46.5 HCl, 1 L-ascorbic acid, 0.5 CaCl2, 5 MgCl2. The slices were incubated in cutting solution at 34°C for 13 minutes, and then maintained at room temperature in extracellular solution until recording, containing (in mM): 124 NaCl, 2.5 KCl, 1.25 NaH2PO4, 26 NaHCO3, 20 glucose, 2 CaCl2, 1 MgCl2.

For recording, the slices were superfused with extracellular solution at 33-35°C. Neurons were visualized using a 60x objective (Olympus, Tokyo, Japan) and a DIC microscope (Scientifica, Uckfield, UK). Patch pipettes (resistance 7-10 MΩ), pulled using a horizontal puller (P-87 Sutter Instruments, Novato, CA, USA) were filled with Cs-internal solution containing (in mM): 130 CsMeSO3, 4 QX-314, 5 Na2-phosphocreatine, 1.5 MgCl2, 10 HEPES, 5 Mg-ATP, 0.35 Na-GTP, 1 EGTA, 8 biocytin for voltage clamp and with 130mM K-gluconate, 5mM KCl, 10mM HEPES, 10mM Na2-phosphocreatine, 4mM ATP-Mg, 0.3mM GTP-Na internal solution for current-clamp recordings. Signals were recorded with an Axon MultiClamp 700B amplifier and digitized at 20 kHz with an Axon Digidata 1550B digitizer (Molecular Devices, San Jose, CA, USA). Access resistance and pipette capacitance were compensated for. In voltage clamp experiments we bath applied TTX (1uM; Tocris) and 4-AP (5 mM; Sigma-Aldrich) to isolate monosynaptic responses. During optogenetic stimulation protocols in all experiments the membrane potential was first depolarized to -40 mV to detect both excitatory and inhibitory currents, then to 0 mV to isolate inhibitory currents which got pharmacologically blocked by bath application GABAA antagonist Gabazine (10uM; Sigma-Aldrich). Then cells were hyperpolarized to -80 mV to isolate AMPA currents which were then blocked by NBQX (20uM; Tocris) and finally the cells got depolarized to +40 mV to measure NMDA-mediated currents which were blocked by the selective NMDA antagonist AP-V (50 uM; Tocris). The synaptic properties of GPi Vglut2+/Vgat+ projections onto GPe neurons were probed using multiple light pulse train protocols with 3 ms blue light pulses with ~2.5 mW light power from a pE-4000 (CoolLED, Andover, UK) light source. To assess passive and active membrane properties, neurons recorded in current-clamp mode were held at a membrane potential of -70 mV. The rheobase current was determined by applying near-threshold current steps, followed by steps proportional to the rheobase current with a duration of 1 s. Intrinsic electrical properties, EPSC and IPSC amplitudes were extracted using a custom-written Matlab (MathWorks, Natick, MA, USA) script.

**Supplementary Table 1 | Rabies experiments cell counts.**

| **Mouse line** | **Injection Site** | **No. Starter Cells** | **Rabies-expressing neurons GPi** | **Animal No.** |
|---|---|---|---|---|
| A2A-cre | CPu | 1240 | 0 | 1 |
| A2A-cre | CPu | 1055 | 0 | 2 |
| A2A-cre | CPu | 1048 | 0 | 3 |
| A2A-cre | CPu | 388 | 0 | 4 |
| OPRm1-cre | CPu | 125 | 8 | 1 |
| OPRm1-cre | CPu | 80 | 5 | 2 |
| OPRm1-cre | CPu | 133 | 6 | 3 |
| OPRm1-cre | CPu | 31 | 8 | 4 |
| OPRm1-cre | CPu | 31 | 4 | 5 |
| OPRm1-cre | CPu | 48 | 3 | 6 |
| DAT-cre | SNc | 388 | 76 | 1 |
| DAT-cre | SNc | 692 | 92 | 2 |
| DAT-cre | SNc | 232 | 23 | 3 |

**Supplementary Table 2 | Mice used in behavioral experiments.**

Behavioral experiments were conducted in the following order: probabilistic two-choice task → arrow maze → open field test → fear conditioning (subset of fiber photometry animals only).

| Experimental group | Mouse ID | Recorded hemisphere | Serial reversal task | Single-reversal maze task | Open field | Fear conditioning |
|---|---|---|---|---|---|---|
| Pv-Cre FIP | 4178 | right | | x<br>session #11 missing FIP | | |
| Pv-Cre FIP | 5036 | right | | x | x | |
| Pv-Cre FIP | 6145 | right | | x<br>#11 missing FIP,<br>#19 & #20 missing | | |
| Pv-Cre FIP | 6881 | right | x | | x | x |
| Pv-Cre FIP | 6892 | right | x | | x | x |
| Pv-Cre FIP | 6914 | right | x | | x | x |
| Sst-Cre FIP | 0520 | left | | x | | |
| Sst-Cre FIP | 6376 | right | | x | x | |
| Sst-Cre FIP | 7061 | left | | x | x | |
| Sst-Cre FIP | 642757 | right | x | x<br>#6, #7, #9 missing FIP | x | |
| Sst-Cre FIP | 651810 | right | x | x<br>#6, #7 missing FIP | | |
| Sst-Cre FIP | 1944 | left | x | | x | x |
| Sst-Cre FIP | 2133 | left | x | | x | x |

| | | | | | | |
|---|---|---|---|---|---|---|
| Sst-Cre FIP | 2144 | left | x | | x | x |
| Sst-Cre FIP | 0419 | left | | | x | |
| Sst-Cre FIP | 6996 | left | | | x | |
| Sst-Cre dTom | 676555 | - | | x | x | |
| Sst-Cre dTom | 676557 | - | | x | x | |
| Sst-Cre dTom | 676558 | - | | x | x | |
| Sst-Cre dTom | 677138 | - | | x | x | |
| Sst-Cre dTom | 705740 | - | | x | x | |
| Sst-Cre dTom | 705742 | - | | x | x | |
| Sst-Cre dTom | 719658 | - | | x | x | |
| Sst-Cre dTom | 719659 | - | | x | x | |
| Sst-Cre dTom | 719660 | - | | x | x | |
| Sst-Cre dTom | 719662 | | | x | | |
| Sst-Cre dTom | 628612 | | | x | | |
| Sst-Cre dTom | 639011 | | | x | | |
| Sst-Cre dTom | 639012 | | | x | | |
| Sst-Cre dTom | 639044 | | | x | | |
| Sst-Cre dTom | 639046 | | | x | | |
| Sst-Cre dTom | 642759 | | | x | | |
| Sst-Cre TeTxLC | 347797 | - | | x | x | |
| Sst-Cre TeTxLC | 347799 | - | | x | x | |

| | | | | | |
|---|---|---|---|---|---|
| Sst-Cre TeTxLC | 347800 | - | | x | x |
| Sst-Cre TeTxLC | 347802 | - | | x | x |
| Sst-Cre TeTxLC | 377803 | - | | x | x |
| Sst-Cre TeTxLC | 347805 | - | | x | x |
| Sst-Cre TeTxLC | 676556 | - | | x | x |
| Sst-Cre TeTxLC | 677132 | - | | x | x |
| Sst-Cre TeTxLC | 677133 | - | | x | x |
| Sst-Cre TeTxLC | 677137 | - | | x | x |
| Sst-Cre TeTxLC | 719097 | - | | x | x |
| Sst-Cre TeTxLC | 719098 | - | | x | x |
| Sst-Cre TeTxLC | 733604 | - | | x | x |
| Sst-Cre TeTxLC | 735151 | - | | x | x |
| Sst-Cre TeTxLC | 735153 | - | | Not learning, interrupted | x |
| Sst-Cre TeTxLC | 736537 | - | | x | x |

### Supplementary Table 3 | Statistical tests.

| Figure | Test | n | Test statistic | P |
|---|---|---|---|---|
| 3C | 6 paired t-tests followed by Holm-Sidak correction | N = 8 | t(7) = [-0.533, -1.093, -2.237, -2.158, -2.105, -1.817] | p = [0.610, 0.311, 0.060, 0.068, 0.073, 0.112]<br>corrected p = [0.610, 0.525, 0.312, 0.312, 0.312, 0.312] |
| 3E | paired t-test | N = 8 | t(7) = 1.112 | p = 0.303 |
| 3F | 6 paired t-tests followed by Holm-Sidak correction | N = 4 | t(3) = [-2.004, 7.782, 4.834, 3.462, 5.900, 13.935] | p = [0.139, 0.004, 0.017, 0.041, 0.010, 0.001]<br>corrected p = [0.139, 0.022, 0.0498, 0.080, 0.038, 0.005] |
| 3H | paired t-test | N = 4 | t(3) = -3.600 | p = 0.037 |
| 3M | Spearman’s rank correlation test (non-parametric) | N = 45 | R = 0.64 | p < 0.001 |
| 3O - Mean speed | Mann-Whitney U test | N = 9 (dTom), N = 16 (TeTxLC) | W = 35 | p = 0.03879 |
| 3O - Angular speed | Mann-Whitney U test | N = 9 (dTom), N = 16 (TeTxLC) | W = 23 | p = 0.004037 |
| 3O - Time immobile | Mann-Whitney U test | N = 9 (dTom), N = 16 (TeTxLC) | W = 123 | p = 0.00325 |
| 3P | Multiple t tests followed by Holm-Sidak correction | N = 9 (dTom), N = 16 (TeTxLC) | df = 1035 | when p < 0.05, a "*" is placed in the figure |
| 4G | repeated-measures ANOVA, within-subject factors: previous outcome, current outcome, and port, including all interaction terms | N = 5 | F(1,4) = [4.649, 228.131, 35.188, 1.011, 1.291, 0.974, 0.174] | p = [0.097, 0.000, 0.004, 0.372, 0.319, 0.380, 0.698] |
| 4N | repeated-measures ANOVA, within-subject factors: previous outcome, current outcome, and port, including all interaction terms | N = 3 | F(1,2) = [3.802, 33.239, 1.002, 0.686, 1.529, 0.005, 1.331] | p = [0.191, 0.029, 0.422, 0.495, 0.342, 0.951, 0.368] |
| 4J | paired t-test | N = 5 | t(4) = 5.779 | p = 0.004 |
| 4K Top | 3 paired t-tests (PE v outcome, PE v % in port, outcome v % in port) followed by Holm-Sidak correction, per port (ipsi, contra) | N = 5 | t(4) = [-4.970, -4.192, 0.461]<br>t(4) = [-1.758, -1.754, 0.536] | p = [0.008, 0.014, 0.669]; corrected p = [0.023, 0.027, 0.669]<br>p = [0.154, 0.154, 0.620]; corrected p = [0.394, 0.394, 0.620] |
| 4K Bottom | 3 t-tests (PE, outcome, % in port; H0: loss = 0) followed by Holm-Sidak correction, per port (ipsi, contra) | N = 5 | t(4) = [-1.885, -2.589, -1.947]<br>t(4) = [-1.703, -0.376, -3.478] | p = [0.133, 0.061, 0.123]; corrected p = [0.232, 0.171, 0.232]<br>p = [0.164, 0.726, 0.025]; corrected p = [0.301, 0.726, 0.074] |
| 4Q | paired t-test | N = 3 | t(2) = 0.480 | p = 0.678 |
| 4R Top | 3 paired t-tests (PE v outcome, PE v % in port, outcome v % in port) followed by Holm-Sidak correction, per port (ipsi, contra) | N = 3 | t(2) = [-7.294, -3.310, 0.601]<br>t(2) = [-1.493, -1.562, -0.506] | p = [0.018, 0.080, 0.609]; corrected p = [0.054, 0.154, 0.609]<br>p = [0.274, 0.259, 0.663]; corrected p = [0.593, 0.593, 0.663] |
| 4R Bottom | 3 t-tests (PE, outcome, % in port; H0: loss = 0) followed by Holm-Sidak correction, per port (ipsi, contra) | N = 3 | t(2) = [-3.730, -2.905, -1.919]<br>t(2) = [-1.306, -0.651, -1.419] | p = [0.065, 0.101, 0.195]; corrected p = [0.182, 0.192, 0.195]<br>p = [0.321, 0.582, 0.292]; corrected p = [0.645, 0.645, 0.645] |
| 5D | 6 paired t-tests followed by Holm-Sidak correction | N = 5 | t(4) = [-1.466, -1.582, -2.283, -4.789, -4.473, -4.915] | p = [0.217, 0.189, 0.084, 0.009, 0.011, 0.008]<br>corrected p = [0.342, 0.342, 0.233, 0.047, 0.047, 0.047] |
| 5E | paired t-test | N = 5 | t(4) = 13.886 | p = 0.000 |

| | | | | |
|---|---|---|---|---|
| 5F | 6 paired t-tests followed by Holm-Sidak correction | N = 3 | t(2) = [-1.911, 2.371, 3.881, 2.313, 3.250, 4.124] | p = [0.196, 0.141, 0.060, 0.147, 0.083, 0.054]<br>corrected p = [0.366, 0.366, 0.284, 0.366, 0.293, 0.284] |
| 5G | paired t-test | N = 3 | t(2) = -23.079 | p = 0.002 |
| 5I - Days 1-14 | Spearman's rank correlation test (non-parametric) | N = 49 | R = -0.15 | p = 0.3 |
| 5I - Days 15-20 | Spearman's rank correlation test (non-parametric) | N = 49 | R = 0.018 | p = 0.9 |
| 5M | Multiple Mann-Whitney U tests followed by Holm-Sidak correction | N = 16 (dTom), N = 15 (TeTxLC) | N/A | when p < 0.05, a "*" is placed in the figure |
| 5N | Multiple Mann-Whitney U tests followed by Holm-Sidak correction | N = 16 (dTom), N = 15 (TeTxLC) | N/A | when p < 0.05, a "*" is placed in the figure |
| 5P | Pearson correlation test | N = 15 | R per each day are reported in the figure | p-values per each day are reported in the figure |
| 6D & E | Kolmogorov-Smirnov test on choice run F in bins "5-14" and "17-20", and on initiation run F in bin "17-20", followed by Holm-Sidak correction | n = [105/1559, 674/88, 61/369] | D = [0.500, 0.203, 0.410] | p = [3.81e-23, 2.70e-03, 1.79e-08]corrected p = [1.14e-22, 2.70e-03, 3.59e-08] |
| 6F | repeated-measures ANOVA | N = 5 | F(3,12) = 20.666 | p = 0.000 |
| 6G | Kolmogorov-Smirnov test on choice run F in bins "5-14" and "17-20", followed by Holm-Sidak correction | n = [75/841, 345/37] | D = [0.172, 0.144] | p = [0.029, 0.446]<br>corrected p = [0.058, 0.446] |
| Sup. Figure 8A | Wald test of the main effect of the slope; linear mixed-effects model with random slopes and intercepts per animal | N = 3 (Pv-Cre), N = 3 (Sst-Cre) | z = [-8.565, -5.058] | p = [1.1e-17, 4.2e-07] |
| Sup. Figure 9C | 8 paired t-tests followed by Holm-Sidak correction | N = 5 | t(4) = [2.061, -0.114, 1.389, -1.112, -1.500, -1.325, -1.006, -0.944] | p = [0.108, 0.915, 0.237, 0.329, 0.208, 0.256, 0.371, 0.398]<br>corrected p = [0.600, 0.915, 0.805, 0.805, 0.805, 0.805, 0.805, 0.805] |
| Sup. Figure 9E | 8 paired t-tests followed by Holm-Sidak correction | N = 3 | t(2) = [0.517, 2.273, 1.596, 1.640, 2.208, 1.189, 1.352, 1.337] | p = [0.657, 0.151, 0.252, 0.243, 0.158, 0.356, 0.309, 0.313]<br>corrected p = [0.811, 0.730, 0.811, 0.811, 0.730, 0.811, 0.811, 0.811] |
| Sup. Figure 9H | 2 repeated-measures ANOVA (trials/min, %correct) | N = 8 | F(3,21) = 56.586<br>F(3,21) = 16.199 | p = 3.114e-10<br>p = 0.000011 |
| Sup. Figure 11B | Multiple Mann-Whitney U tests followed by Holm-Sidak correction | N = 16 (dTom), N = 15 (TeTxLC) | N/A | when p < 0.05, a "*" is placed in the figure |
| Sup. Figure 11C | Multiple Mann-Whitney U tests followed by Holm-Sidak correction | N = 16 (dTom), N = 15 (TeTxLC) | N/A | when p < 0.05, a "*" is placed in the figure |
| Sup. Figure 11D | Multiple Mann-Whitney U tests followed by Holm-Sidak correction | N = 16 (dTom), N = 15 (TeTxLC) | N/A | when p < 0.05, a "*" is placed in the figure |
| Sup. Figure 11E | Multiple Mann-Whitney U tests followed by Holm-Sidak correction | N = 16 (dTom), N = 15 (TeTxLC) | N/A | when p < 0.05, a "*" is placed in the figure |
| Sup. Figure 12C | repeated-measures ANOVA | N = 3 | F(3,6) = 0.583 | p = 0.648 |
